# Mucin glycans signal through the sensor kinase RetS to inhibit virulence-associated traits in *Pseudomonas aeruginosa*

**DOI:** 10.1101/2020.03.31.018614

**Authors:** Benjamin X. Wang, Kelsey M. Wheeler, Kyle C. Cady, Sylvain Lehoux, Richard D. Cummings, Michael T. Laub, Katharina Ribbeck

**Affiliations:** Department of Biology, Massachusetts Institute of Technology, Cambridge, MA; Department of Biological Engineering, Massachusetts Institute of Technology, Cambridge, MA; Howard Hughes Medical Institute, Massachusetts Institute of Technology, Cambridge, MA; Roche Molecular Systems, Inc., Santa Clara, CA; Department of Surgery, Beth Israel Deaconess Medical Center, Harvard Medical School, National Center for Functional Glycomics, Boston, MA

**Author notes:** These authors contributed equally to this work (Benjamin X. Wang, Kelsey M. Wheeler).

## Abstract

Mucus is a densely populated ecological niche that coats all epithelia, and plays a critical role in protecting the human body from infections. Although traditionally viewed as a physical barrier, emerging evidence suggests that mucus can directly suppress virulence traits in opportunistic pathogens like *Pseudomonas aeruginosa*. However, the molecular mechanisms by which mucus affords this protection are unclear. Here, we show that mucins, and particularly their associated glycans, activate the sensor kinase RetS via its Dismed2 domain in *P. aeruginosa*. We find that this RetS-dependent signaling leads to the direct inhibition of the GacS-GacA two-component system, the activity of which is associated with a chronic infection state. This signaling includes the downregulation of the type VI secretion system (T6SS), and prevents T6SS-dependent bacterial killing by *P. aeruginosa*. Overall, these results shed light on how mucus impacts *P. aeruginosa* behavior in the human host, and may inspire novel approaches for controlling *P. aeruginosa* infections.

## Main text

Mucus, a viscoelastic matrix that coats epithelial surfaces, represents a critical interface for host-microbe interactions in the human body, serving as a home for trillions of our commensal microbes (*1*), while simultaneously acting as the first line of defense against many pathogens (*2, 3*). While mucus has historically been viewed as a simple physical barrier, recent work has suggested that mucins, the gel-forming components of mucus, are potent regulators of microbial behavior that can attenuate virulence in a variety of pathogens without affecting viability (*4*–*6*). For example, exposure of the opportunistic pathogen *Pseudomonas aeruginosa* to mucin glycoproteins triggers the suppression of many virulence pathways including quorum sensing and the type III secretion system (*4*). To date, over 200 distinct glycan structures have been identified on mucins (*7*), and may represent a rich source of potential regulatory signals for *P. aeruginosa* and other microbes. However, the molecular mechanisms by which these host-derived signals impact *P. aeruginosa* behavior are unclear (Fig. 1A).

**Figure 1.**
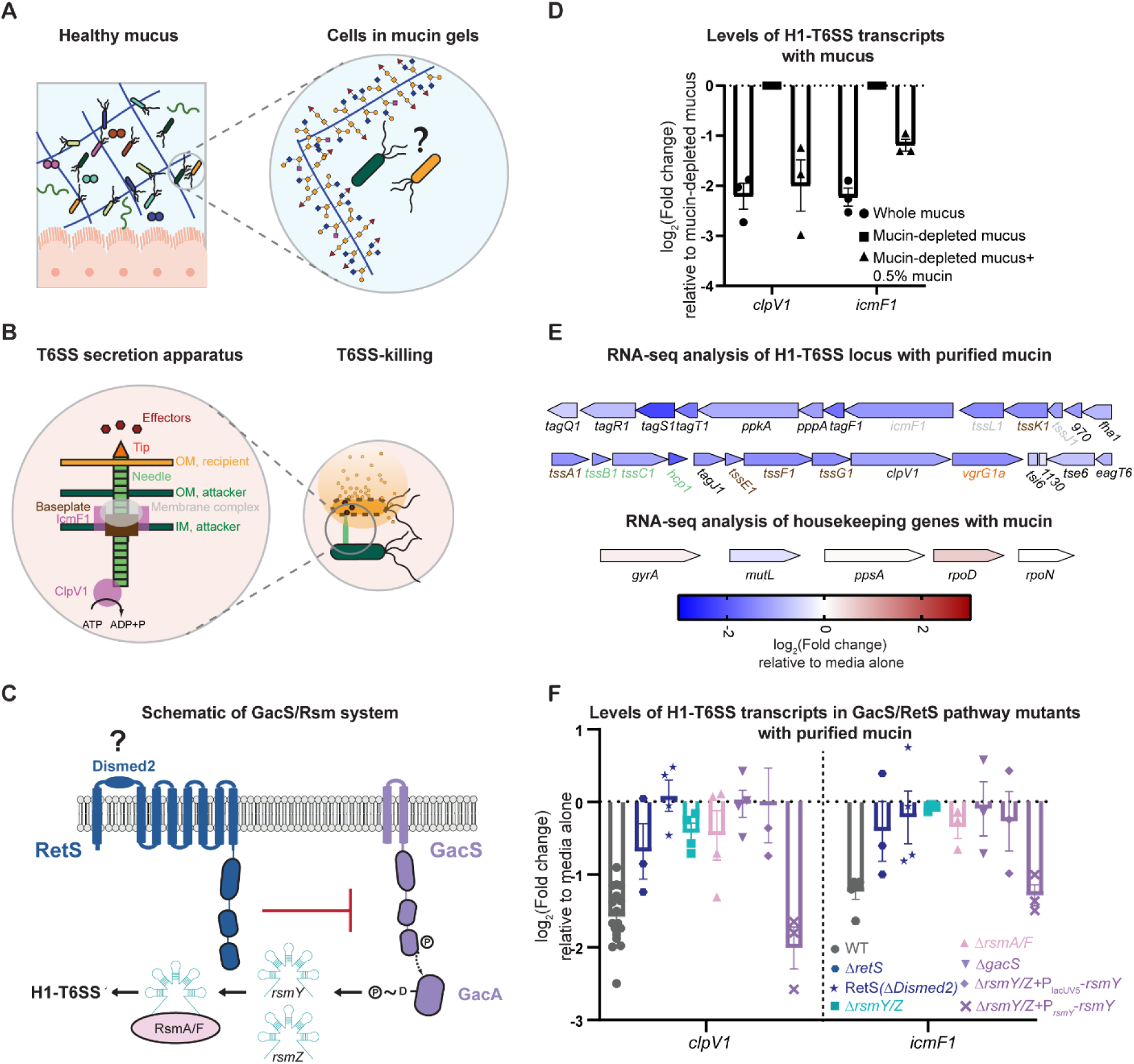
Mucin acts through the GacS/RetS/Rsm pathway to suppress the H1-T6SS. A) Healthy mucus environments suppress bacterial virulence. The mechanisms by which mucin polymers regulate bacterial behavior are unknown. B) Schematic representation of the H1-T6SS needle apparatus. Genes that were measured via qRT-PCR are shown in pink. Other key components of the T6SS are also highlighted: the baseplate in brown, the membrane complex in gray, the needle/sheath in green, the tip in orange, and the effectors in red. C) Schematic representation of the GacS/Rsm system, which controls the expression of the H1-T6SS in response to unknown signals. Upon activation, the histidine kinase GacS autophosphorylates and then transfers its phosphoryl group to its cognate response regulator, GacA. Phosphorylated GacA activates transcription of the regulatory RNAs, *rsmY* and *rsmZ*, which sequester the RNA-binding proteins, RsmA and RsmF. Inhibition of RsmA and RsmF induces changes in the levels of dozens of transcripts, including activation of the H1-T6SS. RetS is an accessory histidine kinase that inhibits GacS. D) Levels of representative H1-T6SS transcripts following 1.5 hour exposure to whole mucus or mucin-depleted mucus with 0.5% MUC5AC added back, relative to mucin-depleted mucus. Transcript levels measured by qRT-PCR and normalized to a control gene (*rpoD*). Bars indicate the mean ± SEM, with individual measurements shown (black dots). E) Diagram of the H1-T6SS operon and housekeeping genes (as a point of reference) representing the fold change of each gene following exposure to mucins for 5 hours relative to medium alone, measured by RNA-sequencing. Arrows indicate the orientation and relative size of each gene. The names of genes that encode key components of the T6SS apparatus are color coded according to Fig. 1B (baseplate in brown, membrane complex in gray, the needle/sheath in green, tip in orange). F) Levels of T6SS transcripts in PA14 wildtype and GacS/Rsm/RetS mutants following exposure to mucin relative to medium alone. Gene expression measured by qRT-PCR and normalized to a control gene (*rpoD*). Bars indicate the mean ± SEM, with individual measurements shown.

One possibility is that *P. aeruginosa* may sense mucin glycans via one of its many two-component systems. Two-component systems are typically comprised of a histidine kinase that senses an environmental signal and a cognate response regulator that triggers changes in gene expression (*8*). Although the signals that activate these two-component systems in *P. aeruginosa* are largely unknown, the histidine kinase RetS may respond to sugars (*9*), making it a candidate for sensing mucin glycans. RetS has been extensively studied for its role in inhibiting GacS-GacA two-component system (*10, 11*), a master regulator of virulence that induces gene expression changes associated with a chronic infection state (*12*). Among these GacS-activated pathways is the Hcp Secretion Island I-encoded Type VI Secretion System (H1-T6SS) (*12*), a needle-like apparatus that directly injects proteinaceous toxic effectors into other bacteria (*13*) (Fig. 1B-C), and may contribute to *P. aeruginosa* fitness during chronic infections (*13, 14*). However, the signals that GacS and RetS respond to have remained elusive.

Here, we identify mucin glycans as signals that activate RetS via its carbohydrate-binding Dismed2 domain to directly inhibit GacS-GacA activity in the *P. aeruginosa* PA14 strain. In particular, we find glycan signaling downregulates the H1-T6SS and suppresses T6SS-dependent bacterial killing by PA14 in a RetS-dependent manner. Collectively, these results provide new insights into the mechanisms by which signals in healthy mucus are sensed by pathogens to suppress virulence-associated behaviors, and may guide the design of new molecules for mitigating infections by problematic pathogens like *P. aeruginosa*.

### Mucins downregulate the type VI secretion system

For *P. aeruginosa*, a chronic infection state is associated with activation of antagonistic factors such as the H1-T6SS (*12, 14*). However, it is unclear how the H1-T6SS is regulated in native healthy mucus, in which *P. aeruginosa* is typically unable to establish infections. To test how healthy mucus impacts the H1-T6SS, we cultured PA14, a hypervirulent clinical isolate, in a solution of whole mucus containing ∼0.5% MUC5AC, an isoform of mucin present in niches colonized by *P. aeruginosa*. We used qRT-PCR to quantify the levels of *clpV1*, an ATPase involved in the secretion of type VI effectors, and *icmF1*, which encodes a structural component of the secretion apparatus (Fig. 1B). The levels of these representative H1-T6SS transcripts, which are present on two different operons in this locus, were both lowered by ∼4-fold in whole mucus (Fig. 1D), relative to mucin-depleted mucus. Furthermore, addition of purified MUC5AC (0.5%) to mucin-depleted mucus restored H1-T6SS downregulation (Fig. 1D), indicating that mucins are sufficient to suppress the H1-T6SS in native mucus. The magnitude of changes (∼3-4 fold) seen here with mucin are similar to changes in H1-T6SS levels seen in response to *P. aeruginosa* lysate (*15*).

In addition to *clpV1* and *icmF1*, the H1-T6SS locus contains 25 other genes. To determine the extent to which mucin regulates the rest of the H1-T6SS locus, we performed RNA-sequencing on PA14 exposed to purified mucins for 5 hours in chemically defined medium (Table S1). Isolated MUC5AC downregulated H1-T6SS genes by an average of 2.3-fold, with 18 of the 27 transcripts downregulated at least 2-fold (Fig. 1E, Fig. S1A). For comparison, five housekeeping genes were not substantially affected (>|2-fold|) by mucins (Fig. 1E). In agreement with the RNA-seq data, purified mucins decreased *clpV1* and *icmF1* transcript levels approximately 2-3 fold when measured by qRT-PCR (Fig. S1B), which was similar to the downregulation observed with whole mucus (Fig. 1D). Taken together, these results suggest that mucins downregulate the majority of H1-T6SS genes in PA14.

### Mucins activate the sensor kinase RetS, resulting in the inhibition of GacS and the downstream Rsm pathway

RetS is a sensor kinase that inhibits the H1-T6SS (*12*) (Fig. 1C). This protein contains an N-terminal periplasmic Dismed2 domain that has homology to carbohydrate-binding proteins (*9*), and could potentially respond to mucin glycoproteins. To determine if RetS can sense mucins, we generated a strain lacking RetS, and found the ability of mucins to downregulate *clpV1* and *icmF1* in this background was impaired (Fig. 1F). Similarly, a variant of RetS lacking its periplasmic Dismed2 domain, which was as stable as full-length RetS protein (Fig. S1C), also failed to respond to mucin (Fig. 1F).

RetS controls H1-T6SS activity by inhibiting the GacS-GacA two-component system and the downstream Rsm pathway (Fig. 1C). To test if mucins act through the Gac/Rsm pathway, we deleted key components of the pathway, namely GacS (Δ*gacS*), *rsmY/Z (*Δ*rsmY/Z*), and RsmA/RsmF (Δ*rsmA/F*), and exposed each of these strains to mucin. Mucin was unable to downregulate H1-T6SS transcripts in any of these strains (Fig. 1F), suggesting these components are essential to mucin-mediated suppression of the H1-T6SS. However, for the Δ*gacS* and Δ*rsmY/Z* strains, H1-T6SS levels may be low enough (Fig. S1D) that mucin cannot further downregulate c*lpV1* or *icmF1*. To circumvent this limitation, we complemented the Δ*rsmY/Z* strain with a plasmid expressing *rsmY* from the constitutive lacUV5 promoter (Δ*rsmY/Z +* P_lacUV5_-*rsmY*), which restored *clpV1* and *icmF1* to approximately wild-type levels (Fig. S1D) but removed GacS/GacA control of *rsmY* expression. Mucins were unable to downregulate the H1-T6SS in the Δ*rsmY/Z +* P_lacUV5_-*rsmY* strain (Fig. 1F). By contrast, when the plasmid-borne copy of *rsmY* was expressed from its native promoter, mucin could downregulate *clpV1* and *icmF1* (Fig. 1F). Taken together, our results indicate that mucins act through RetS and the Gac/Rsm pathway to downregulate the H1-T6SS.

### Mucin glycans are the mucus-derived signal that mediate the downregulation of H1-T6SS

Given the predicted sugar binding function of the RetS Dismed2 domain, mucin-derived glycans may be the component of mucins that activate RetS. To test this possibility, we isolated mucin glycans from MUC5AC by alkaline β-elimination. Using matrix-assisted laser desorption/ionization time-of-flight (MALDI-TOF) mass spectrometry, we identified >80 glycan structures by mass (Fig. 2A, Table S2), which does not include possible isomeric forms. This diversity of glycan structures in the purified pool offers a rich source of potential signals that may activate RetS. Indeed, exposure to 0.1% w/v of this mucin glycan pool (corresponding to low millimolar concentrations) was sufficient to downregulate *clpV1* and *icmF1* (Fig. 2B). We also measured T6SS transcript levels following exposure to a pool of 0.1% monosaccharides representing those present in mucin. The monosaccharide pool did not suppress *clpV1* or *icmF1* (Fig. 2B), suggesting that the complex structures of mucin glycans are critical to their function.

**Figure 2.**
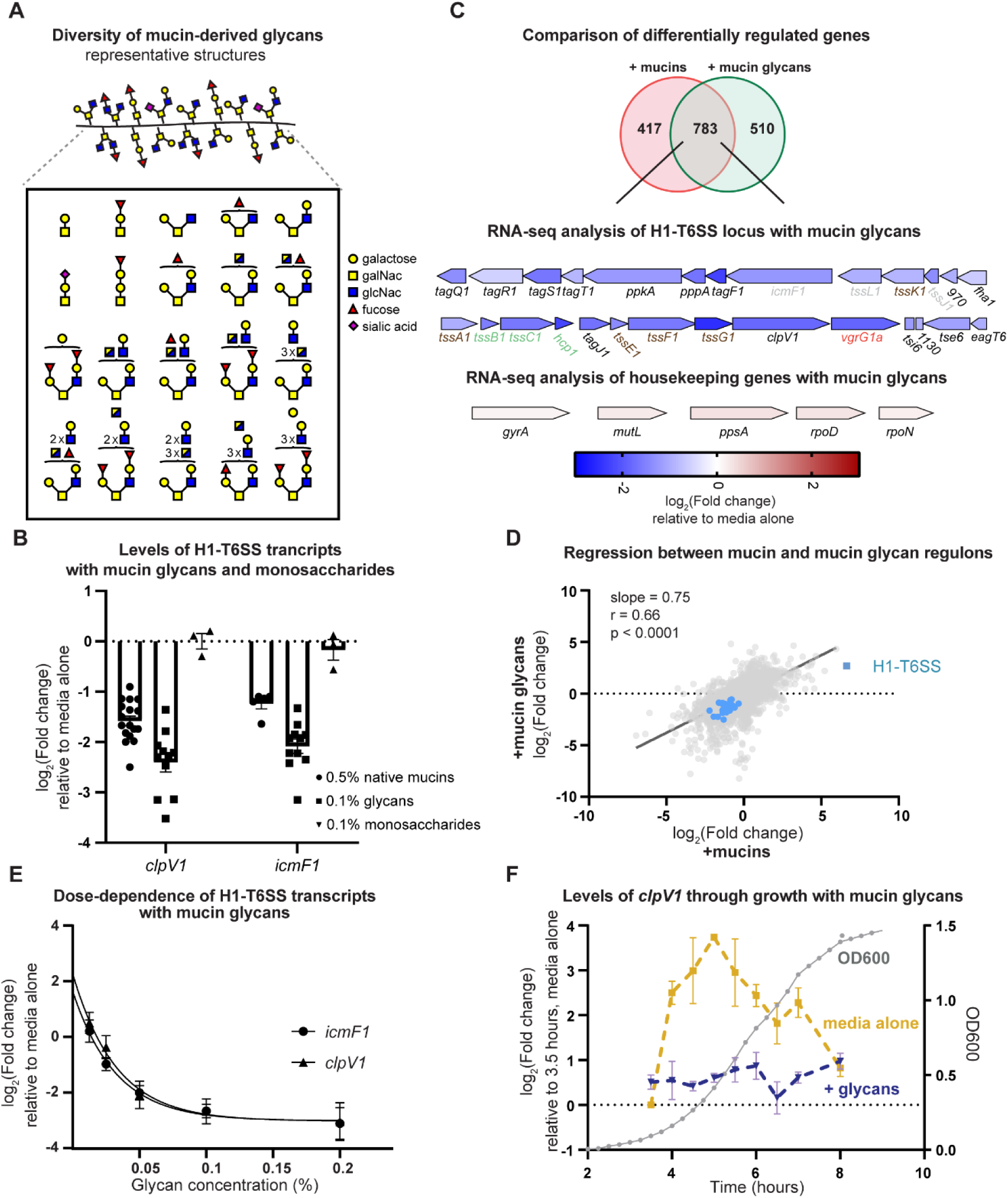
Mucin glycans are sufficient to suppress the T6SS. A) Representative glycan structures derived from MUC5AC, as identified by mass spectrometry. B) Levels of T6SS transcripts in PA14 following exposure to the pool of mucin glycans or monosaccharides for 5 hours relative to medium alone. Gene expression measured by qRT-PCR and normalized to a control gene (*rpoD*). Bars indicate the mean ± SEM, with individual measurements shown (black dots). Mucin data are replicated from Fig. 1F, for reference. C) Overlap of differentially expressed genes in response to mucins or glycans after 5 hours relative to medium alone, measured by RNA-sequencing. Diagram of the H1-T6SS operon and housekeeping genes representing the fold change of each gene following exposure to glycans for 5 hours relative to medium alone, measured by RNA-sequencing. Arrows indicate the orientation and relative size of each gene. The names of genes that encode key components of the T6SS apparatus are color coded according to Fig. 1B (baseplate in brown, membrane complex in gray, the needle/sheath in green, tip in orange). D) Regression of glycan-induced transcriptional changes against mucin-induced transcriptional changes at 5 hours. Genome wide transcriptional changes were measured by RNA-sequencing. H1-T6SS genes are highlighted in blue. E) Levels of T6SS transcripts in PA14 following 5-hour exposure to the pool of mucin glycans at different concentrations relative to medium alone. Gene expression measured by qRT-PCR and normalized to a control gene (*rpoD*). Data points indicate the mean ± SEM and are fitted to a one-site binding curve. F) Levels of T6SS transcripts in PA14 following exposure to the pool of mucin glycans (in blue) or medium alone (in yellow) at different time points, as measured by qRT-PCR and normalized to a control gene (*rpoD*). Transcript levels are measured relative to medium alone at 3.5 hours, and each point represents the mean ± SEM (3 replicates). Grey data points indicate optical density of the culture at 600 nm.

To determine the extent to which free mucin glycans recapitulate the potency of whole intact mucins, we used RNA-seq to quantify the effects of 0.1% glycans on the full H1-T6SS locus (Table S1). Similar to the suppression of H1-T6SS seen with intact mucins (Fig. 1E), H1-T6SS genes were downregulated by an average of 2.6-fold in the presence of glycans (Fig. 2C, Fig. S2A). Beyond the H1-T6SS, there was also a significant correlation (r = 0.66, p < 0.0001) between the gene expression profiles in response to mucin and glycans, indicating that glycans are largely responsible for the gene expression changes induced by mucin (Fig. 2D).

To further characterize the response to mucin glycans, we measured *clpV1* and *icmF1* levels at multiple glycan concentrations, and found that serially diluted glycans downregulated these transcripts in a dose-dependent manner (Fig. 2E). Further, glycans suppressed *clpV1* expression at multiple time points throughout exponential phase (Fig. 2F), suggesting that the glycan response is prolonged and not an artifact of a particular time point.

Signaling through RetS and GacS was previously found to trigger gene expression changes within 15-25 minutes (*15*). To determine how quickly the H1-T6SS is downregulated in response to mucin glycans, we performed RNA-seq on cells exposed to glycans for 15 minutes (Fig. S2B) following growth to mid-exponential phase, when the H1-T6SS is maximally expressed in the absence of glycans (Fig. 2F). We found that H1-T6SS locus was downregulated by an average of ∼2-fold after 15 minutes of glycan exposure (Fig. S2C, Table S1). To confirm these RNA-seq results, we used qRT-PCR to measure *clpV1* and *icmF1* levels after incubation with glycans for 15 minutes, and also observed a ∼2-fold downregulation (Fig. S2D). These results indicate that mucin glycans rapidly downregulate the H1-T6SS.

### Mucin glycans activate RetS via its periplasmic Dismed2 and its cytoplasmic phosphorelay domains

To confirm that liberated mucin glycans, like their intact mucin counterparts, also act through GacS and RetS, we incubated glycans with the Δ*gacS*, Δ*retS*, and signal-blind RetS(Δ*Dismed2*) strains, and measured H1-T6SS transcripts with qRT-PCR. As with intact mucins, free mucin glycans were no longer able to downregulate *clpV1* or *icmF1* in any of these mutants (Fig. 3A), confirming that both the sensory domain of RetS and GacS are necessary for glycan signaling.

**Figure 3.**
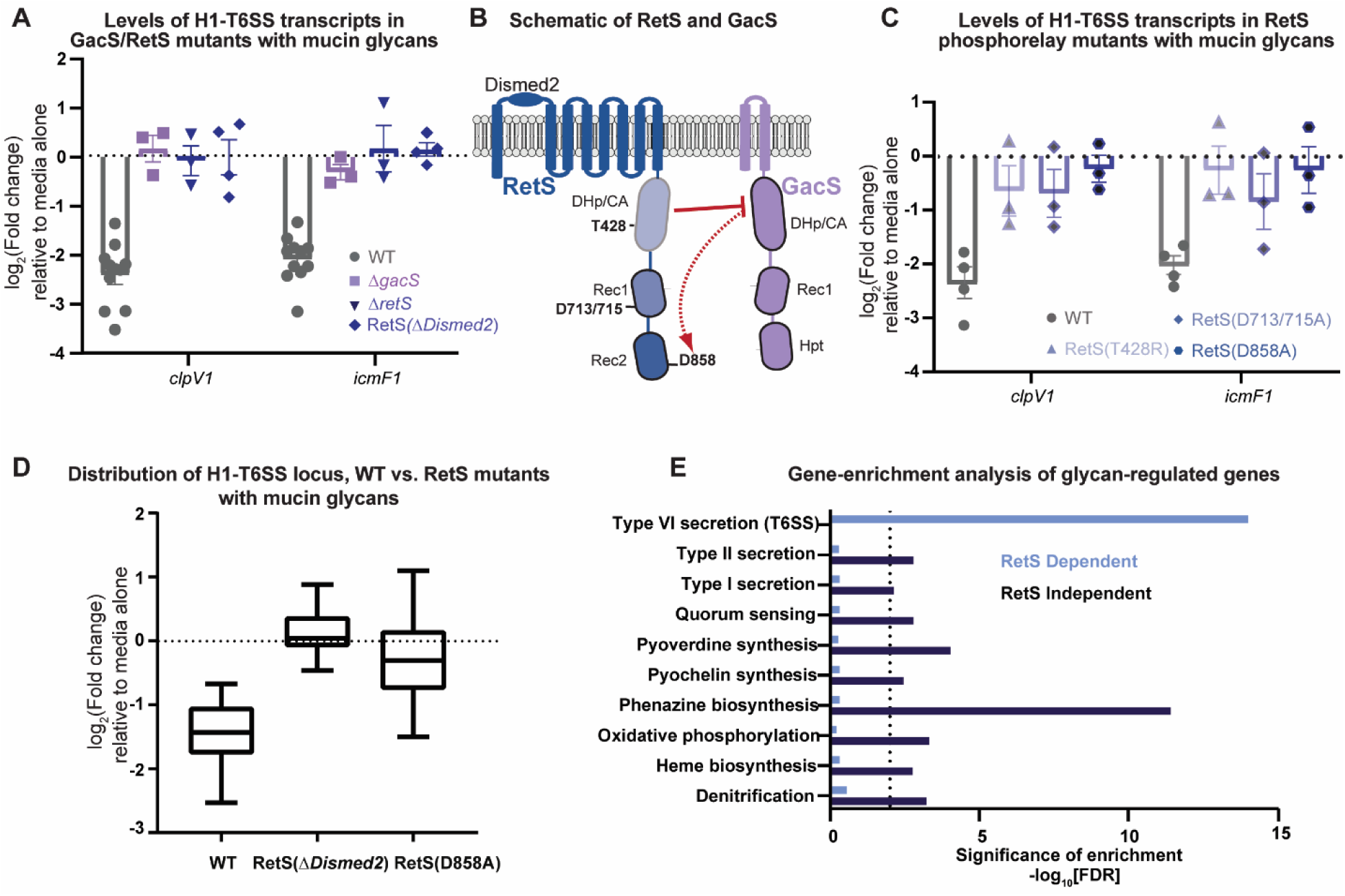
Mucin glycans promote RetS inhibition of GacS through the RetS phosphorelay domains. A) Levels of T6SS transcripts in PA14 RetS, RetS(Δ*Dismed2*) and GacS mutants following exposure to mucin glycans for 5 hours relative to medium alone. Gene expression measured by qRT-PCR and normalized to a control gene (*rpoD*). Bars indicate the mean ± SEM, with individual measurements shown. Wild type data are duplicated from Fig. 2B for reference. B) Schematic of the phosphorelay domains of RetS (various shades of blue) and GacS/GacA (purple), with key phosphorelay residues numbered and domains labeled.RetS likely inhibits GacS both by acting as a phosphatase via its DHp/CA domain and by siphoning phosphates from GacS onto its second receiver domain. C) Levels of T6SS transcripts in RetS phospho-mutants following exposure to mucin glycans for 5 hours relative to medium alone. Gene expression measured by qRT-PCR and normalized to a control gene (*rpoD*). Bars indicate the mean ± SEM, with individual measurements shown. D) Distribution of all H1-T6SS transcript levels following exposure to mucin glycans for 5 hours relative to a medium control in the wild type and RetS mutants, as measured by RNA-sequencing. Center bar represents the mean change in gene expression for all genes in the T6SS operon. E) Gene set enrichment analysis of RetS-dependent (i.e., glycan-mediated changes that only occur in the wild type) and RetS-independent (i.e., glycan-mediated changes that occur in the wild type and both RetS mutants) gene sets. Bar length indicates false discovery rate for each enriched pathway. The dotted line indicates the threshold for significance.

Although these results indicate that mucin glycans inhibit the GacS pathway by activating RetS, it is unclear how this signaling event is conveyed through these two histidine kinases at a molecular level. Because GacS activity depends on its phosphorylation state, we investigated the role of the three phosphorelay domains in RetS, which were described to modulate GacS activity in other *P. aeruginosa* strains to varying degrees (*16*–*18*), but have not been investigated in PA14. Specifically, we mutated a conserved residue of the DHp/CA domain to ablate potential phosphatase activity (T428R), and conserved residues in the two tandem receiver domains to prevent RetS from siphoning phosphoryl groups from GacS (D713/715A, D858A) (Fig. 3B). After confirming that these mutations did not affect RetS stability (Fig. S3A), we measured GacS activity in these mutants by using qRT-PCR to quantify H1-T6SS transcript levels (Fig. S3B), and by using an *rsmY-lacZ* reporter (Fig. S3C). These assays revealed that both the DHp/CA and second receiver domains of RetS are critical for GacS inhibition in PA14. Furthermore, the ability of RetS(D858) to receive phosphate from GacS was confirmed using *in vitro* phosphorylation assays (Fig. S3D).

Having confirmed the roles of the RetS phosphorelay domains in PA14, we then tested the involvement of these domains in glycan signaling. We found that the ability of glycans to suppress *clpV1* and *icmF1* expression was impaired in all three RetS point mutants to varying degrees, with the RetS(D858A) mutant displaying the strongest loss of glycan-induced T6SS suppression (Fig. 3C). Overall, these results are consistent with a model in which glycans simulate RetS to directly siphon phosphate from GacS and to act as a phosphatase, which leads to decreased GacS activity and suppression of the H1-T6SS.

### Mucin glycan signaling through RetS affects the RetS regulon

To confirm that glycans act through RetS to suppress the rest of the H1-T6SS locus in addition to *clpV1* and *icmF1*, we performed RNA-seq on the signal-blind RetS(Δ*Dismed2*) and RetS(D858A) strains following exposure to mucin glycans (Fig. S4A-B, Table S1). While glycans downregulated the H1-T6SS genes by ∼2.6-fold on average in the wild type, this locus was no longer downregulated by glycans in either RetS mutant (Fig. 3D, Fig. S4C).

RetS controls the expression of dozens of other genes beyond the H1-T6SS, many of which are uncharacterized but may contribute to chronic infections (*12*). To determine the extent to which these other RetS-dependent genes are regulated by mucin glycans, we compared gene expression profiles of glycan-exposed wild-type cells to the published expression profile of the Δ*retS* strain (*12*) (Table S3). These two profiles were inversely correlated (r = −0.57, p<0.0001), as glycans inhibit GacS, whereas a deletion of *retS* activates GacS (Fig. S4D). By contrast, there was no significant correlation when comparing the gene expression profiles of glycan-exposed RetS(Δ*Dismed2*) and RetS(D858A) cells to the published Δ*retS* regulon (Fig. S4E).

A previous study reported that mucins regulate other pathways in addition to the H1-T6SS, many of which are virulence-associated (*4*). To determine if these pathways were also suppressed via RetS, we performed gene set enrichment analysis on the glycan-regulated genes of the wild-type, RetS(Δ*Dismed2*), and RetS(D858A) strains. Of all the pathways significantly enriched in the wild-type strain following glycan exposure, the H1-T6SS was the only pathway not enriched in either RetS mutant (Fig. 3E, Fig. S5). By contrast, all other pathways were differentially expressed in the wild-type and both RetS mutants (Fig. 3E, Fig. S5).

Overall, these findings demonstrate that glycans act specifically through RetS to suppress H1-T6SS and the majority of the RetS regulon, but likely act through different pathways to control RetS-independent phenotypes.

### *P. aeruginosa* likely senses mucin glycans built on a core 2 structure

Mucin glycans are built on either a linear core 1 (Galβ1-3GalNAcα1-) or branched core 2 (Galβ1-3(GlcNAcβ1-6)(GalNAcα1-) structure. Often, these glycans are further elongated with additional galactose residues that are modified with terminal sialic acid and/or fucose. To determine if these terminal moieties are essential for glycan signaling, we applied partial acid hydrolysis to the glycan pool. Sialic acid is removed by hydrolysis at 0.1 M trifluoroacetic acid (TFA) and fucose is removed by 1 M TFA, while the core structures are resistant to acid hydrolysis (*19*).

We identified putative structures of the 1M TFA-treated glycans by mass spectrometry (Fig. 4A-B, Table S2). Acid treatment generated shorter glycans, with the average number of monosaccharides per glycan decreasing from 4.46 in the untreated pool to 2.65 in the acid-treated sample (Fig. 4C). Glycans in both the untreated and acid-treated pools retained high levels of the acid-resistant core 1 and core 2 structures (Fig. 4D). By contrast, acid-treated glycans had substantially less fucose and sialic acid compared to the untreated pool (Fig. 4E).

**Figure 4.**
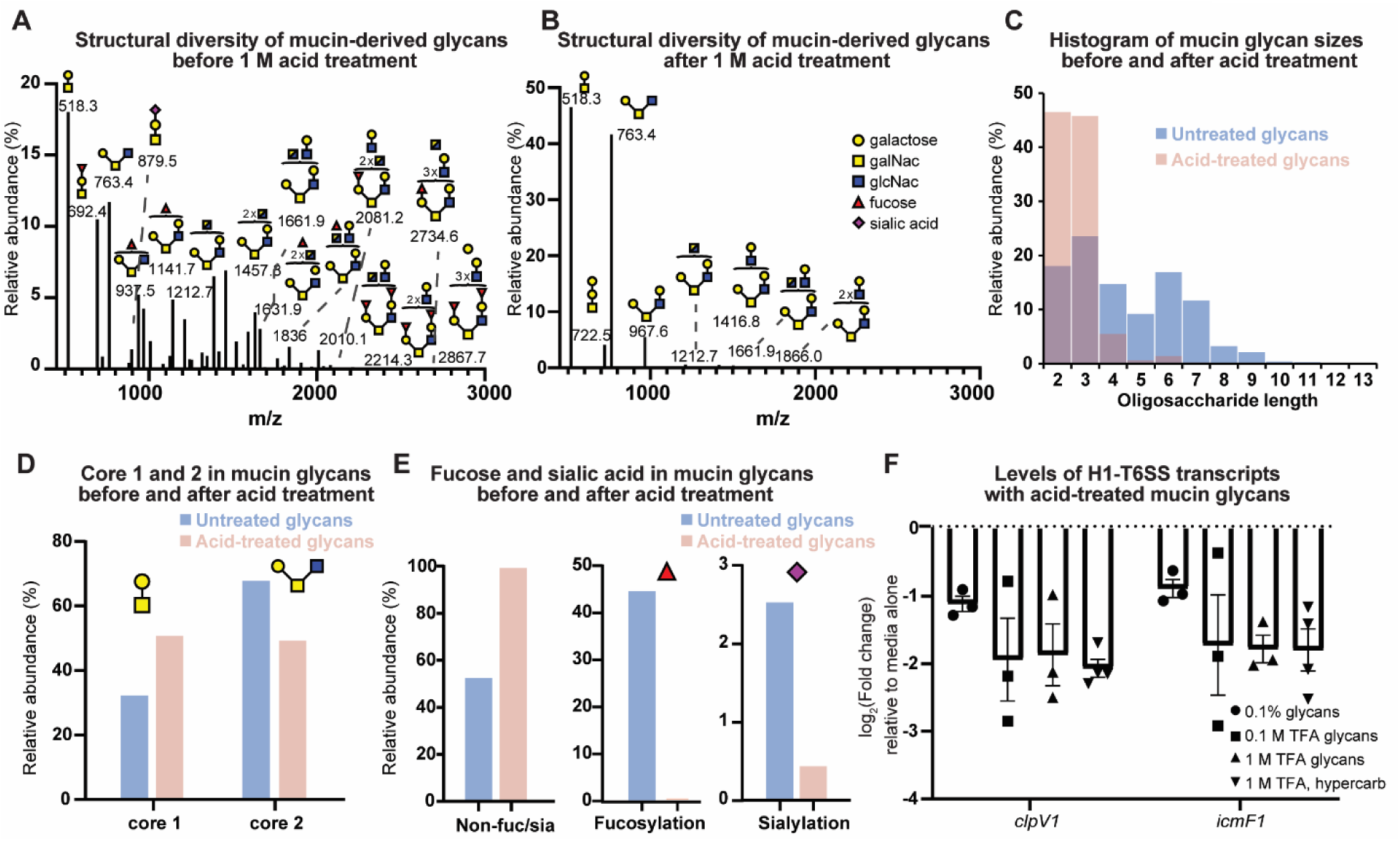
Active glycans likely contain a core 2 moiety, but do not require fucose or sialic acid. A) MALDI-TOF-MS spectrum of the full MUC5AC glycan pool. Selected peaks labeled with monoisotopic masses and predicted structures. B) MALDI-TOF-MS spectrum of the MUC5AC glycan pool following partial acid hydrolysis. Selected peaks labeled with monoisotopic masses and predicted structures. C) Distribution of glycan length in the untreated versus the acid-treated glycan pools. D) Relative abundance of core structures in the full pool of MUC5AC glycans versus the pool of glycans following acid treatment. E) Relative abundance of fucosylated, sialylated, and non-fucosylated/sialylated glycans in the full MUC5AC glycan pool versus the acid-treated glycan pool. F) Levels of T6SS transcripts in PA14 following exposure to pools of acid-treated glycans for 5 hours relative to medium alone. Gene expression measured by qRT-PCR and normalized to a control gene (*rpoD*). Bars indicate the mean ± SEM, with individual measurements shown (black dots).

Notably, acid-treated glycans were at least as potent as the diverse pool, triggering a strong downregulation of *clpV1* and *icmF1* (Fig. 4F). Thus, it is likely that glycans do not require fucosylation or sialylation to function as a signal. To confirm that the active signal is an intact glycan, and not an acid-released monosaccharide, we fractionated the 1 M TFA-treated glycans through a Hypercarb cartridge, which retains intact glycans but not monosaccharides or salts (*20*). The composition of the glycan pool before and after Hypercarb purification was similar, suggesting that most intact glycans were successfully retained by the column (Fig. S6A, Table S2). These Hypercarb-retained glycans downregulated *clpV1* and *icmF1* to the same extent as the unseparated glycans (Fig. 4F). We further tested commercially-available sugar moieties present in mucin glycans including the core 1 structure, N-Acetyl-D-lactosamine (lacNAc), and the oligosaccharides (GlcNAc)_2_ and (GlcNAc)_3_. None of these di- or trisaccharides downregulated the H1-T6SS (Fig. S6B), although the possibility remains that natively purified glycans may differ from their synthesized counterparts in various ways including linkage, modifications, or the presence of associated molecules. Overall, these results indicate that H1-T6SS suppressing glycans are likely built on a core 2 structure and do not require terminal fucose or sialic acid.

### Mucin glycans suppress T6SS-mediated killing by PA14 through RetS

Downregulation of the H1-T6SS by mucin glycans should lead to decreased killing of neighboring bacteria by PA14. To test this prediction, we used an established T6SS killing assay involving the co-incubation of PA14 and *E. coli* (*15*). In the absence of glycans, co-incubation of PA14 and *E. coli* led to substantial killing of *E. coli* (∼1-2 logs) (Fig. 5A). Deleting a critical structural component of the H1-T6SS (Δ*icmF1*) abrogated killing (Fig. 5A), confirming that *E. coli* killing was dependent on the H1-T6SS under these experimental conditions. Strikingly, the addition of glycans also abolished *E. coli* killing to a similar extent as the Δ*icmF1* mutant (Fig. 5A). To determine if this loss of killing was dependent on the T6SS or occurred through an independent pathway, we repeated the competition between *E. coli* and the Δ*icmF1* strain in the presence of glycans, and found that glycans did not further increase *E. coli* survival in this competition (Fig. 5A).

**Figure 5.**
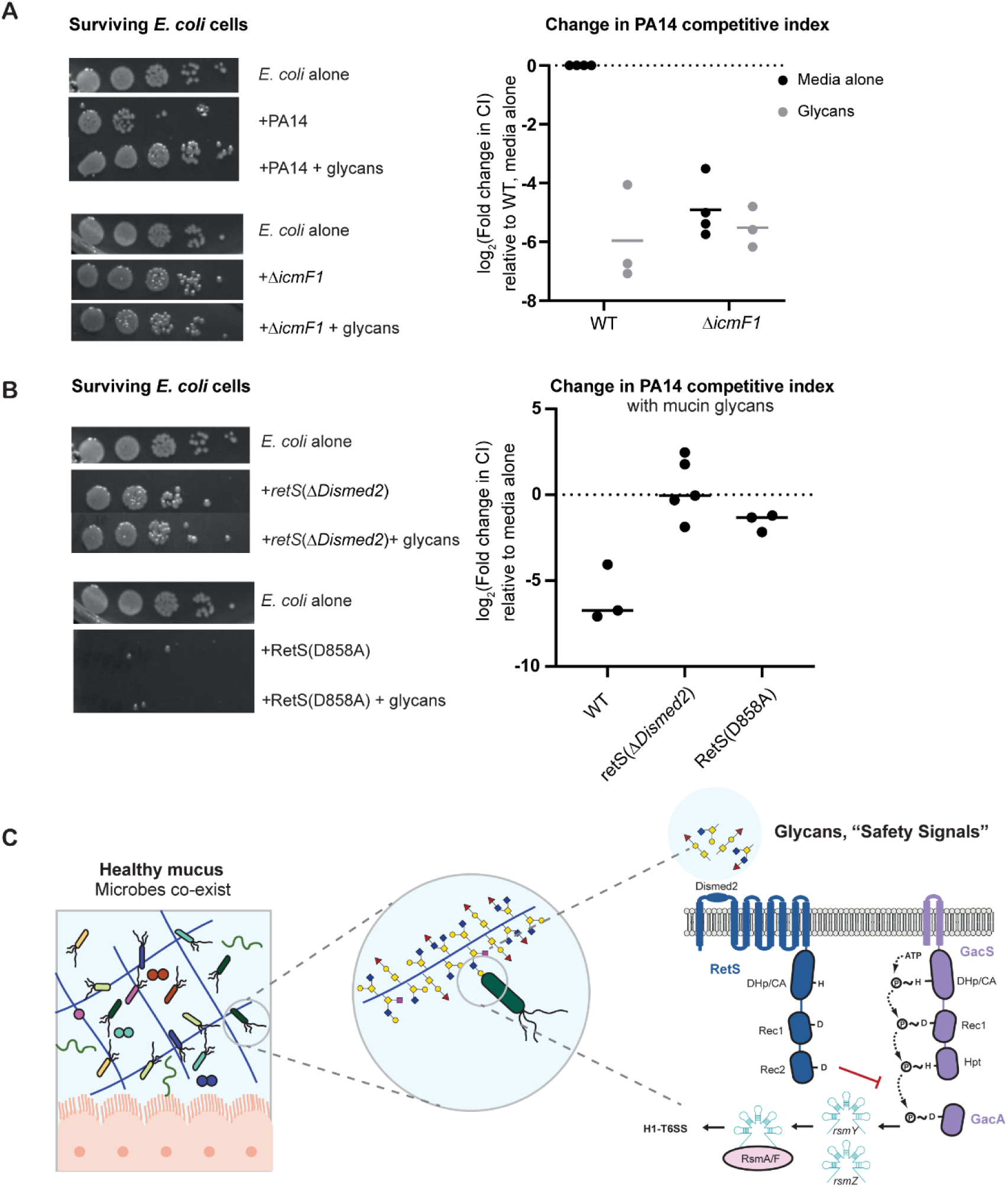
Mucin glycans suppress T6SS-mediated killing by PA14 in a RetS-dependent manner. A) Competition assay between *E. coli* and the PA14 wild type and T6SS mutant (Δ*icmF1*) in the presence and absence of glycans. Left: Representative images of serial dilution of viable *E. coli* cells following competition with PA14. Right: Center bar represents the mean change in *P. aeruginosa* competitive index relative to the wild type in medium alone, with individual measurements shown (black dots). B) Competition assay between *E. coli* and the PA14 RetS mutants in the presence and absence of glycans. Left: Representative images of serial dilution of viable *E. coli* cells following competition with PA14. Right: Center bar represents the mean change in *P. aeruginosa* competitive index in the presence of glycans relative to medium alone, with individual measurements shown (black dots). C) Mucin glycans are host-produced “safety signals” that suppress the H1-T6SS by activating RetS via its carbohydrate-binding Dismed2 sensory domain, which directly inhibits GacS activity. The resulting suppression of the H1-T6SS may help prevent *P. aeruginosa* from colonizing healthy mucus.

Because H1-T6SS downregulation by glycans was abolished in both the RetS(Δ*Dismed2*) and RetS(D858A) mutants (Fig. 3D, Fig. S4C), we reasoned that glycans would no longer be able to rescue T6SS-mediated killing of *E. coli* by PA14 in these mutant strains. To test this hypothesis, we performed killing assays between *E. coli* and these PA14 mutants in the presence or absence of glycans. The RetS(D858A) mutation leads to hyper-activation of the T6SS (Fig. S3B-C), and accordingly we observed a drastic loss in *E. coli* viability compared to competitions with wild-type PA14 (Fig. 5B). Notably, glycans were no longer able to rescue *E. coli* survival when co-cultured with either RetS mutant strain (Fig. 5B), confirming that glycans act through both the Dismed2 and second receiver domains of RetS to suppress T6SS-mediated killing.

## Discussion

Recent work indicates that mucus contains potent signals that influence microbial gene expression and behavior (*4*–*6*). However, the molecules in mucus and the mechanisms by which bacteria sense and respond to these signals have remained unclear. Here, we demonstrate that a subset of mucin glycans activates the sensor kinase RetS via its Dismed2 sensory domain in *P. aeruginosa*. This signaling stimulates the ability of RetS to inhibit GacS, likely by siphoning phosphoryl groups from GacS and by acting as a phosphatase. Mucin glycans induce changes in the expression of the majority of the GacS regulon including the downregulation of the H1-T6SS, which prevents T6SS-dependent killing of *E. coli* by *P. aeruginosa* in a RetS-dependent manner.

While RetS likely responds to a subset of mucin glycans built on a core 2 structure, some homologs of Dismed2 can bind to multiple glycans (*21*–*24*). Thus, other carbohydrates may also signal through RetS. For example, upon lysis, *P. aeruginosa* is thought to release a “danger signal” that signals through RetS to activate the H1-T6SS in kin cells (*15*). Although the exact signal was not found, an intriguing hypothesis is that the signal may be a bacterially-produced carbohydrate that resembles mucin glycans, and that specific structural differences between these bacterial and host signals may either activate or inhibit RetS. In this way, perhaps mucin glycans can be considered a host-produced “safety signal” that suppresses bacterial antagonism and instead promotes microbial co-existence (Fig. 5C). Indeed, our observation that mucin glycans lead to reduced T6SS-dependent killing in phenotypic assays may help explain the inability of *P. aeruginosa* to invade and displace communities found in healthy mucus.

We anticipate that changes to mucin glycosylation in certain diseases may prevent the virulence-suppressing response documented here. In particular, increased sialyation and distinct glycosylation patterns have been observed in the mucus of individuals with cystic fibrosis (*25*–*27*), which may partly explain how *P. aeruginosa* is capable of establishing infections in this niche. Further dissection of how various glycans influence RetS activity may inform the development of intervention strategies to limit *P. aeruginosa* infections.

The ability of mucin glycans to activate a bacterial receptor suggests that the host provides a rich source of signals that can be perceived by microbes. Consistent with this hypothesis, mucin glycans built on the core 2 structure prevent pathogenic *E. coli* invasion into a colonic cell line through an unknown mechanism (*28*), suggesting that the virulence-attenuating effects of core 2-containing glycans are not exclusive to *P. aeruginosa*. Furthermore, purified mucins attenuate virulence in the Gram-positive *Streptococcus mutans* (*5*) and the fungal pathogen *Candida albicans* (*6*), but specific mucin-sensing receptors have not been identified in these species. Ultimately, the large assortment of receptors across these phylogenetically distant species, combined with the rich diversity of mucin glycans, underscores the enormous signaling potential that is housed within mucus. An understanding of the molecular mechanisms driving these signaling events may inspire the design of novel therapeutics that can attenuate virulence in a variety of problematic pathogens.

## Acknowledgements

The authors thank M. LeRoux, M. Guzzo, K. Gozzi, and B. Imperiali for comments on the manuscript. This research was supported by NIBIB/NIH grant R01 EB017755-04 (OSP 6940725) (to K.R.), the MRSEC Program of the National Science Foundation under award DMR-1419807 (to K.R.), the National Science Foundation Career award PHY-1454673 (to K.R.), a CEHS grant (P30-ES002109) (to K.R.), the National Science Foundation Graduate Research Fellowship under grant no. 1745302 (to B.X.W. and K.M.W.), the National Center for Functional Glycomics Grant P41GM103694 (to R.D.C.), and an NIH grant to M.T.L. (R01GM082899), who is also an Investigator of the Howard Hughes Medical Institute.

## Author contributions

B.X.W., K.M.W, K.R., and M.T.L. designed the experiments. K.C. performed preliminary experiments related to the RetS-GacS phosphorelay and constructed strains. B.X.W. and K.M.W. performed all experiments and analyses. S.L. and R.D.C. performed the glycan mass spectrometry. B.X.W., K.M.W., K.R., and M.T.L. wrote the paper.

## Data and materials availability

Raw RNA-sequencing data will be made available on GEO.

## Supplementary Materials

Materials and Methods

Figures S1-S6

Tables S1-S7

References (*29-35*)

## References

1. P. C. Barko, M. A. McMichael, K. S. Swanson, D. A. Williams, The Gastrointestinal Microbiome: A Review. J. Vet. Intern. Med. 32, 9–25 (2018).

2. M. Zanin, P. Baviskar, R. Webster, R. Webby, The Interaction between Respiratory Pathogens and Mucus. Cell Host Microbe. 19 (2016), pp. 159–168.

3. T. Pelaseyed, J. H. Bergström, J. K. Gustafsson, A. Ermund, G. M. H. Birchenough, A. Schütte, S. van der Post, F. Svensson, A. M. Rodríguez-Piñeiro, E. E. L. Nyström, C. Wising, M. E. V. Johansson, G. C. Hansson, The mucus and mucins of the goblet cells and enterocytes provide the first defense line of the gastrointestinal tract and interact with the immune system. Immunol. Rev. 260, 8–20 (2014).

4. K. M. Wheeler, G. Cárcamo-Oyarce, B. S. Turner, S. Dellos-Nolan, J. Y. Co, S. Lehoux, R. D. Cummings, D. J. Wozniak, K. Ribbeck, Mucin glycans attenuate the virulence of Pseudomonas aeruginosa in infection. Nat. Microbiol. (2019), doi:10.1038/s41564-019-0581-8.

5. E. S. Frenkel, K. Ribbeck, Salivary mucins protect surfaces from colonization by cariogenic bacteria. Appl. Environ. Microbiol. 81, 332–338 (2015).

6. N. L. Kavanaugh, A. Q. Zhang, C. J. Nobile, A. D. Johnson, K. Ribbeck, Mucins suppress virulence traits of Candida albicans. MBio. 5 (2014), doi:10.1128/mBio.01911-14.

7. C. Jin, D. T. Kenny, E. C. Skoog, M. Padra, B. Adamczyk, V. Vitizeva, A. Thorell, V. Venkatakrishnan, S. K. Lindén, N. G. Karlsson, Structural diversity of human gastric mucin glycans. Mol. Cell. Proteomics. 16, 743–758 (2017).

8. E. J. Capra, M. T. Laub, Evolution of Two-Component Signal Transduction Systems. Annu. Rev. Microbiol. 66, 325–347 (2012).

9. V. Anantharaman, L. Aravind, Application of comparative genomics in the identification and analysis of novel families of membrane-associated receptors in bacteria. BMC Genomics. 4 (2003), doi:10.1186/1471-2164-4-34.

10. S. Heeb, D. Haas, “Regulatory Roles of the GacS/GacA Two-Component System in Plant-Associated and Other Gram-Negative Bacteria” (2001).

11. K. Lapouge, M. Schubert, F. H. T. Allain, D. Haas, Gac/Rsm signal transduction pathway of γ-proteobacteria: From RNA recognition to regulation of social behaviour. Mol. Microbiol. 67 (2008), pp. 241–253.

12. A. L. Goodman, B. Kulasekara, A. Rietsch, D. Boyd, R. S. Smith, S. Lory, A signaling network reciprocally regulates genes associated with acute infection and chronic persistence in Pseudomonas aeruginosa. Dev. Cell. 7, 745–754 (2004).

13. R. D. Hood, P. Singh, F. S. Hsu, T. Güvener, M. A. Carl, R. R. S. Trinidad, J. M. Silverman, B. B. Ohlson, K. G. Hicks, R. L. Plemel, M. Li, S. Schwarz, W. Y. Wang, A. J. Merz, D. R. Goodlett, J. D. Mougous, A Type VI Secretion System of Pseudomonas aeruginosa Targets a Toxin to Bacteria. Cell Host Microbe. 7, 25–37 (2010).

14. J. D. Mougous, M. E. Cuff, S. Raunser, A. Shen, M. Zhou, C. A. Gifford, A. L. Goodman, G. Joachimiak, C. L. Ordoñez, S. Lory, T. Walz, A. Joachimiak, J. J. Mekalanos, A virulence locus of Pseudomonas aeruginosa encodes a protein secretion apparatus. Science (80-.). 312, 1526–1530 (2006).

15. M. Le Roux, R. L. Kirkpatrick, E. I. Montauti, B. Q. Tran, S. Brook Peterson, B. N. Harding, J. C. Whitney, A. B. Russell, B. Traxler, Y. A. Goo, D. R. Goodlett, P. A. Wiggins, J. D. Mougous, Kin cell lysis is a danger signal that activates antibacterial pathways of pseudomonas aeruginosa. Elife. 2015, 1–65 (2015).

16. M. A. Laskowski, B. I. Kazmierczak, Mutational analysis of RetS, an unusual sensor kinase-response regulator hybrid required for Pseudomonas aeruginosa virulence. Infect. Immun. 74, 4462–4473 (2006).

17. V. I. Francis, E. M. Waters, S. E. Finton-James, A. Gori, A. Kadioglu, A. R. Brown, S. L. Porter, Multiple communication mechanisms between sensor kinases are crucial for virulence in Pseudomonas aeruginosa. Nat. Commun. 9 (2018), doi:10.1038/s41467-018-04640-8.

18. A. L. Goodman, M. Merighi, M. Hyodo, I. Ventre, A. Filloux, S. Lory, Direct interaction between sensor kinase proteins mediates acute and chronic disease phenotypes in a bacterial pathogen. Genes Dev. 23, 249–259 (2009).

19. A. Guttman, Analysis of monosaccharide composition by capillary electrophoresis. J. Chromatogr. A. 763, 271–277 (1997).

20. N. H. Packer, M. A. Lawson, D. R. Jardine, J. W. Redmond, A general approach to desalting oligosaccharides released from glycoproteins. Glycoconj. J. 15, 737–747 (1998).

21. E. Fouquaert, E. J. M. van Damme, Promiscuity of the Euonymus carbohydrate-binding domain. Biomolecules. 2 (2012), pp. 415–434.

22. A. L. van Bueren, A. B. Boraston, The Structural Basis of α-Glucan Recognition by a Family 41 Carbohydrate-binding Module from Thermotoga maritima. J. Mol. Biol. 365, 555–560 (2007).

23. A. B. Boraston, V. Notenboom, R. A. J. Warren, D. G. Kilburn, D. R. Rose, G. Davies, Structure and ligand binding of carbohydrate-binding module CsCBM6-3 reveals similarities with fucose-specific lectins and “galactose-binding” domains. J. Mol. Biol. 327, 659–669 (2003).

24. E. Ficko-Blean, A. B. Boraston, N-Acetylglucosamine Recognition by a Family 32 Carbohydrate-Binding Module from Clostridium perfringens NagH. J. Mol. Biol. 390, 208–220 (2009).

25. B. Xia, J. A. Royall, G. Damera, G. P. Sachdev, R. D. Cummings, Altered O-glycosylation and sulfation of airway mucins associated with cystic fibrosis. Glycobiology. 15, 747–775 (2005).

26. T. F. Scanlin, M. C. Glick, Terminal glycosylation in cystic fibrosis. Biochim. Biophys. Acta. 1455, 241–53 (1999).

27. M. Davril, S. Degroote, P. Humbert, C. Galabert, V. Dumur, J. J. Lafitte, G. Lamblin, P. Roussel, The sialylation of bronchial mucins secreted by patients suffering from cystic fibrosis or from chronic bronchitis is related to the severity of airway infection. Glycobiology. 9, 311–321 (1999).

28. J. Ye, Q. Pan, Y. Shang, X. Wei, Z. Peng, W. Chen, L. Chen, R. Wang, Core 2 mucin-type O-glycan inhibits EPEC or EHEC O157:H7 invasion into HT-29 epithelial cells. Gut Pathog. 7, 1–9 (2015).

29. T. Masuko, A. Minami, N. Iwasaki, T. Majima, S. I. Nishimura, Y. C. Lee, Carbohydrate analysis by a phenol-sulfuric acid method in microplate format. Anal. Biochem. 339, 69–72 (2005).

30. P. H. Culviner, C. K. Guegler, M. T. Laub, bioRxiv, in press, doi:10.1101/2020.01.06.896837.

31. T. T. Hoang, A. J. Kutchma, A. Becher, H. P. Schweizer, Integration-proficient plasmids for Pseudomonas aeruginosa: Site-specific integration and use for engineering of reporter and expression strains. Plasmid. 43, 59–72 (2000).

32. A. Brencic, S. Lory, Determination of the regulon and identification of novel mRNA targets of *Pseudomonas aeruginosa* RsmA. Mol. Microbiol. 72, 612–632 (2009).

33. S. Castang, S. L. Dove, Basis for the essentiality of H-NS family members in Pseudomonas aeruginosa. J. Bacteriol. 194, 5101–5109 (2012).

34. R. M. Q. Shanks, N. C. Caiazza, S. M. Hinsa, C. M. Toutain, G. A. O’Toole, Saccharomyces cerevisiae-based molecular tool kit for manipulation of genes from gramnegative bacteria. Appl. Environ. Microbiol. 72, 5027–5036 (2006).

35. K. A. McFarland, E. L. Dolben, M. LeRoux, T. K. Kambara, K. M. Ramsey, R. L. Kirkpatrick, J. D. Mougous, D. A. Hogan, S. L. Dove, A self-lysis pathway that enhances the virulence of a pathogenic bacterium. Proc. Natl. Acad. Sci. U. S. A. 112, 8433–8438 (2015).

