## Supplemental materials for "Mucin glycans signal through the sensor kinase RetS to inhibit virulence-associated traits in *Pseudomonas aeruginosa*"

**This PDF file includes:**

Materials and Methods

Figs. S1 to S6

Tables S4 to S7

**Other Supplementary Materials for this manuscript include the following:**

Tables S1 to S3 titles

### Materials and Methods

#### General cloning procedures

*P. aeruginosa* and *E. coli* strains were both grown overnight in LB at 37 $^{o}$C under shaking conditions. When applicable, antibiotics were added at the following concentrations: for *E. coli,* 10 µg/mL tetracycline, 20 µg/mL gentamycin, and 100 µg/mL carbenicillin, and for *P. aeruginosa,* 100 µg/mL tetracycline, and 75 µg/mL gentamycin. Strains, plasmids, and primers used are listed in Supplemental Tables 4-7.

Chromosomal-based modifications in *P. aeruginosa* (deletions, insertions, and point mutations) were done with allelic exchange using the suicide vectors pEXG2 and pMQ30, which both confer gentamycin resistance and sucrose sensitivity. In general, vectors were made by Gibson assembly (N.E.B) and transformed into S17 *E. coli* on LB with appropriate antibiotics. S17 strains were then grown up to $OD_{600}$ ~ 0.6. At the same time, the relevant *P. aeruginosa* strains were incubated at 42 $^{o}$C for at least 2 hours. Then, 1.5 mL of S17 and 0.5 mL of *P. aeruginosa* were mixed, spun down, and resuspended in 100 µL LB. This mixture was then pipetted onto the center of a plain LB plate and grown overnight at 30 $^{o}$C for mating to occur. The mixture was then isolated and re-suspended in 1 mL PBS, with dilutions of this solution plated on *Pseudomonas* isolation agar (PIA) supplemented with the appropriate antibiotics. Single colonies were then streaked on LB plates supplemented with 15% sucrose for counter selection. Colonies from the sucrose plates were then patched onto plain LB plates and subjected to colony PCR; clones with the correct constructs were collected and stored at -80 $^{o}$C.

#### Plasmid and reporter construction

The pEXG2 allelic exchange vector was used to generate the Rsm deletion strains, the *icmF1* mutant, as well as the RetS(Δ*Dismed2*) chromosomal construct. In this case, fragments corresponding to regions ~500-1000 bases upstream and downstream from each deleted sequence were PCR amplified from PA14 gDNA. Primers were designed so that the upstream and downstream fragments had ~20-30 bp overlap regions, and so that each fragment also overlapped by ~20-30 bp with the pEXG2 vector. Next, fragments were cloned into the pEXG2 vector via Gibson assembly (N.E.B) using overlapping regions between the two fragments, and between each fragment and the vector. After sequence confirmation, assembled vectors were electroporated into S17 cells and mated with WT PA14. Colony PCR was performed on single colonies from the LB+15% sucrose plates, and those with the correct deletions were re-streaked onto plain LB plates. Complementation of the *rsmY/Z* deletion was done by expressing *rsmY* from either a constitutively active lacUV5 promoter or its native promoter on the pPSV38plasmid. After sequence confirmation, the pPSV38-*rsmY* plasmid was electroporated into S17 cells and subjected to the same procedures described above, with the *rsmY/Z* deletion used during mating.

Deletions of *retS*, *gacS,* and *hptB* were made by allelic exchange using the suicide vector pMQ30. Briefly, regions corresponding to ~500-1000 bases upstream and downstream of these genes were amplified from PA14 gDNA. Primers were designed so that the two amplified fragments had ~20-30 bp of overlap, and so that both fragments had ~20-30 bp of overlap with the pMQ30 vector. Fragments were then assembled with linearized pMQ30 in yeast. Sequence verified plasmids were then transformed into S17 and subjected to the same procedures described above, with wild-type PA14 used during mating.

The pMQ30 allelic exchange vector was also used to generate chromosomal point mutations in *retS*. However, in this case, primers were designed so that the overlapping region between the upstream and downstream fragments contained the relevant point mutations. Fragments were then amplified from PA14 gDNA and combined with linearized pMQ30 in yeast. Subsequent steps were identical to that described above, except that plasmids were conjugated into a Δ*retS* strain to introduce mutant *retS* sequences at the native chromosomal locus.

The pEXG2 suicide vector was also used to add chromosomal 3x-FLAG tag to RetS variants. Briefly, an ~1.3 kb fragment comprising ~650 bases upstream and downstream of the desired 3x-FLAG integration site was amplified from PA14 gDNA and put into pEXG2 by Gibson assembly (N.E.B). Then, around-the-horn PCR was used to add the nucleotide sequence encoding a 3x-FLAG tag at the desired site in the assembled vector by putting each half of the 3x-FLAG tag region on the forward and reverse primers. The rest of the procedure was identical to that described above for construction of RetSΔDISMED2, except that the *P. aeruginosa* strains used for mating were different, depending on the desired background. Colony PCR was performed on individual clones from the LB+15% sucrose plates to check for integration of the 3x-FLAG tag.

For the RetS(D858A)-3xFLAG construct, the D858A mutation introduced was within the region used for recombination. As such, around-the-horn PCR was used to generate the D858A mutation on the assembled pEXG2-RetS-3xFLAG vector by designing a forward primer containing the mutation at the 5' end.

The *rsmY-lacZ* chromosomal reporters were made using the previously published pCTX-*rsmY-lacZ* plasmid. Briefly, the plasmid was transformed into electrocompetent S17 *E. coli* cells and mated with the relevant *P. aeruginosa* strains. Single colonies that appeared on PIA supplemented with tetracycline were then picked, re-streaked on LB-tetracycline plates, and subjected to colony PCR to confirm integration of the *rsmY-lacZ* construct.

Expression vectors for protein purification were made using the pENTR-TOPO gateway cloning kit (Invitrogen). Briefly, $GacS_c$ and $RetS_c$ sequences were amplified from PA14 gDNA, with a 'CACC' sequence added to the 5' end of each forward primer. PCR products were then incubated with the pENTR vector according to the Topo kit instructions, and then transformed into chemically competent DH5α *E. coli* cells. After sequence verification, the correct pENTR vectors were subjected to a LR recombination reaction with either ML333 or ML310 destination vectors using the Gateway LR Clonase II enzyme mix (Invitrogen). Sequence verified plasmids were transformed into BL21 *E. coli* cells for protein expression.

To generate point mutations of RetS for *in vitro* assays, primers were designed so that the mutation of interest was located near the middle of both forward and reverse primers. Mutations were then generated by using a Quikchange site-directed mutagenesis protocol. Sequence verified plasmids were transformed into BL21 *E. coli* cells for protein expression.

### Mucus experiments

Mucus was scraped from fresh pig stomachs and solubilized (1 g scrapings to 5 mL) in 0.2 M sodium chloride buffer with protease inhibitors (5 mM Benzamidine HCl, 1 mM dibromoacetophenone, 1 mM phenylmethylsulfonylfluoride (PMSF), and 5mM EDTA at pH 7) and 0.04% sodium azide (Sigma). Cellular debris and food waste was removed via low-speed centrifugation, 8000 x g, (7,000 rpm Sorvall GS-3 rotor), for 30 minutes at 4 °C. Mucus was depleted of its high molecular weight components with Amicon ultra centrifugal 100 kDa filters (Sigma).

Batch cultivation of *Pseudomonas aeruginosa* PA14 was carried out shaking at 37 °C in LB (Difco). For whole-mucus experiments, overnight cultures of PA14 were diluted 150-fold into ABTGC (ABT minimal medium supplemented with 0.2% glucose, and 0.2% casamino acids) and grown shaking at 37 °C for 3.5 h. Thirty microliters of these cultures were exposed to 30 µL of solubilized mucus, depleted mucus, or solubilization buffer for 1.5 h at 37 °C in a static 96-well microtiter plate (Cellstar).

### Mucin purification

This study used native porcine gastric mucins (MUC5AC). Mucus was scraped from fresh pig stomachs, solubilized in sodium chloride buffer (described above), and insoluble material was removed by ultracentrifugation at 190,000 x g RCF for 1 hr at 4 °C (40,000 rpm, Beckman 50.2 Ti rotor with polycarbonate bottles). Mucus was then clarified and desalted with disposable PD-10 desalting columns (GE). Mucins were isolated using size-exclusion chromatography on a Sepharose CL-2B column. Mucin fractions were identified based on absorbance at 280 nm, then desalted and concentrated with an Amicon stirred cell pressure-based concentrator (Sigma) with an Omega ultrafiltration 100 kDa membrane disc filter (Pall). Purified mucins were then lyophilized for storage at -80 °C.

Lyophilized mucins are reconstituted by shaking them gently at 4 °C overnight in the desired medium.

**Mucin glycan isolation**

This study applied non-reductive alkaline β-elimination ammoniolysis to dissociate non-reduced glycans from commercially available pig gastric mucins (Sigma). Mucins were added to 1x PBS at a ratio of 30 mg per mL and insoluble material was removed by low-speed centrifugation, 8000 x g, (7,000 rpm Sorvall GS-3 rotor), for 20 minutes at 4 °C. Mucins were then precipitated with 60% (v/v) ethanol, collected by centrifugation, and dissolved in water. Dissolved mucin was then desalted and concentrated with an Amicon stirred cell pressure-based concentrator (Sigma) with 100 kDa membrane filter disc (Pall), and lyophilized. Lyophilized mucins were dissolved in ammonium hydroxide saturated with ammonium carbonate and incubated at 60°C for 48 h to release oligosaccharide glycosylamines and partially deglycosylated mucins. Volatile salts were removed via repeated centrifugal evaporation and the oligosaccharide glycosylamines were separated from residual deglycosylated mucins via centrifugal filtration through 10 kDa molecular weight cut-off membranes in accordance with the manufacturer's instructions (Amicon Ultracel). The resulting oligosaccharide glycosylamines were converted to reducing oligosaccharide hemiacetals via treatment with boric acid. Residual boric acid was removed via repeated centrifugal evaporation from methanol.

Glycans were analyzed using matrix-assisted laser desorption/ionization time-of-flight (MALDI-TOF). β-eliminated glycans were permethylated and analyzed at the Glycomics Core at Beth Israel Deaconess Medical Center. Mass spectrometry data were acquired on an UltraFlex II MALDI-TOF Mass Spectrometer (Bruker Daltonics). Reflective positive mode was used, and data were recorded between 500 m/z and 6000 m/z. The mass spectrometry O-glycan profile was acquired by aggregating at least 20,000 laser shots. Mass peaks were manually annotated and assigned to a particular O-glycan composition based on known core structures.

Glycans were quantified using the phenol-sulfuric method (*29*). Briefly, dried glycans were dissolved in 250 µL of water. 25 µL of glycans were added to a 96 well plate, following by

150 µL of concentrated sulfuric acid with rigorous pipetting. 30 µL of a 5% phenol solution was then added, which caused the solution to turn brown. Quantification of this color change was measured at 490 nm, and w/v measurements were made by comparing to a standard curve of serially diluted glucose.

**Acid treatment of mucin glycans**

Mucin glycans were incubated with 1 M TFA at 80 °C for 4 hours to remove sialic acid and fucose. Glycans were then dried using a speedvac, and washed repeatedly with methanol until all TFA was removed, as verified by pH testing.

For samples further purified through Hypercarb cartridges (Thermo), acid-treated glycans were first neutralized with potassium hydroxide (KOH), incubated at room temperature for 10 minutes to allow salts to precipitate, and then spun at 16,000g for 10 minutes to remove any precipitant. The supernatant was then collected and dried using a speedvac.

At the same time, the Hypercarb cartridges were first primed with 4 column volumes of 100% acetonitrile, then flushed with 4 column volumes of water. Dried glycans were resuspended in 500 µL of water, and loaded onto the cartridge. The Hypercarb cartridge was then washed with 2 column volumes of water and 1 column volume of 2% acetonitrile to remove salts and monosaccharides. Glycans were then eluted using 2 column volumes of 50% acetonitrile, and dried using a speedvac.

**Set-up of sugar-signaling experiments**

Cells were grown in ABTGC media (15.1 mM ammonium sulfate, 33.7 mM sodium phosphate dibasic, 22.0 mM potassium dihydrogen phosphate, 0.05 mM sodium chloride, 1 mM magnesium chloride, 100 µM calcium chloride, 1 µM iron (III) chloride, 0.2% glucose, and 0.2% casamino acids). Unless otherwise indicated, cells were grown in 96 well format for 5 hours (~30 µL/well), and reached an OD of ~0.5-0.6.

For mucin experiments, mucin was reconstituted in ABTGC at 0.5% w/v overnight (4 degrees, with constant vortexing). The next morning, overnight cultures were diluted 1:100 into reconstituted mucin solutions and grown at 37 °C for 5 hours. For glycan experiments, dried glycans were reconstituted to 0.1% in ABTGC unless otherwise indicated, and overnight cultures were diluted 1:100 into these solutions and grown at 37 $^{o}$C for 5 hours. For monosaccharide experiments, the monomeric forms of each mucus sugar (GlcNac, GalNac, galactose, fucose, sialic acid), were purchased from Sigma and dissolved as a 0.1% w/v solution in ABTGC, and were then used in the same way as reconstituted mucins and glycans.

**RNA preparation for qRT-PCR and RNA-seq experiments**

30 µL of cells was pelleted at max speed in a table top centrifuge. Cell pellets were then resuspended in 300 µL of Cell Lysis solution (Epicentre) with 2 uL of 50 µg/µL proteinase K and incubated at 65 $^{o}$C for 30 minutes, with vortexing every 5 minutes. Samples were placed on ice for 5 minutes, and then mixed with 175 µL of MPC Protein Precipitation Reagent (Epicentre). Next, samples were spun for 10 minutes at maximum speed in a tabletop centrifuge at 4 $^{o}$C. The supernatant was then transferred to a clean tube and mixed with 500 µL of isopropanol by inverting the tubes ~30 times. Tubes were spun for 10 minutes at maximum speed in a tabletop centrifuge at 4 $^{o}$C again to pellet the RNA. After removal of the supernatant, the RNA pellet was mixed with 0.5 mL of 70% ethanol and centrifuged at 12,000 *g* for 15 minutes at 4 $^{o}$C. The RNA pellet was washed again with 70% ethanol, and then air dried for 10 minutes at room temperature and resuspended in 30 µL of nuclease free water.

To remove contaminating DNA from RNA preps, 5 µL of Ambion RNA Turbo buffer + 1.5 µL of Turbo DNase (Invitrogen) were added to each RNA sample and incubated for 30 minutes at 37 $^{o}$C. Following this incubation, 5 uL of the Ambion reaction inactivator (Invitrogen) were added to each sample and vortexed rigorously. After incubation at room temperature for 2 minutes, samples were spun down for 2 minutes at 10,000 *g* with a tabletop centrifuge. The supernatant containing purified RNA was then carefully collected and transferred to a new tube, before storing at -80 $^{o}$C for long-term storage.

**qRT-PCR**

The SuperScript III kit (Invitrogen) was used to prepare cDNA for qRT-PCR experiments. Briefly, 1 µg of purified RNA was combined with 1 µL of 50 ng/µL random hexamers, 1 μL of a 10 mM dNTP mix, and nuclease free water to a total volume of 10 μL. The samples were incubated at 65 °C for 5 minutes, and then placed on ice for 5 minutes. Next, 2 μL of 10X RT buffer, 4 μL of 25 mM $MgCl_2$, 2 μL of 0.1 M DTT, 1 μL of RNaseOUT™, and 1 μL of SuperScript III were added to each sample, and then incubated at 25 °C for 10 minutes, 50 °C for 50 minutes, and 85 °C for 5 minutes. Samples were then stored at -20 °C for long-term storage or used immediately for qRT-PCR.

qRT-PCR was performed with the Sybr Fast qPCR 2x Master Mix (Kapa). Briefly, ~10 ng of cDNA was incubated with 5μL of 2x Sybr, 0.6 μL of 5 μM of each primer, for a final primer concentration of 300 nM, and nuclease free $H_2O$ to a final volume of 10 μL. Samples were then placed onto 384 well white bottom plates, and qRT-PCR done in a LightCycler 480 system (Roche) with the following thermocycling program: 95 °C for 10 minutes, 95 °C for 15 seconds, 60 °C for 30 seconds, and 72 °C for 30 seconds, with 40 cycles of steps 2-4.

Data were analyzed by the ddCT method to determine the fold change of each relative to a housekeeping gene (*rpoD*). Primers were checked for efficiency by running serially diluted concentrations of cDNA, and only primers with efficiency of >90% were used.

**RNA-seq**

Purified RNA was depleted of rRNA using biotinylated rRNA-specific probes (*30*). Following ribo-depletion, 15 μL of depleted RNA was submitted to the MIT BioMicro center for strand-specific library preparation. Single-end sequencing was performed on a HiSeq2000, with a read length of 40 nucleotides. BWA was used to map the reads to the PA14 genome, and uniquely mapping reads were used. Any read that overlapped with a gene region was given a count of 1 read.

The RNA-seq data was processed as follows. Genes with an average of fewer than 10 counts were eliminated, as well as all rRNA reads. Next, the total number of reads across all genes was calculated for both the + and – glycan conditions. The normalized read count for each gene was calculated by dividing each raw read count by the total read count for each condition. Finally, the fold change of each gene was calculated as the ratio between the normalized read count of each gene in the + vs. –condition (ie, +/- mucins, +/- glycans), for each respective strain and/or time point.

RNA-seq data will be made available in GEO.

**Western blotting**

For Western blotting experiments, 50 mL of cells were grown up to $OD_{600}$ ~ 0.8, and then spun down and washed three times in 50 mM $KH_2PO_4$ at pH 7.5. Cells were then resuspended in 1 mL of 50 mM $KH_2PO_4$ and lysed by sonication (2 minutes total time, 1 second on, 4 second off cycles at 30% power). Lysates were spun down for 10 minutes at 5,000 *g* to remove unlysed cells. As GacS and RetS are membrane proteins, proteins were not denatured by boiling, which can induce oligomerization of membrane proteins. Instead, lysates were incubated with 4x SB (200 mM Tris-HCl pH 6.8, 8% SDS, 0.4% bromophenol blue, 40% glycerol) for 30 minutes at room temperature to gently denature proteins and then sonicated in a Diagenode water bath sonicator for 5 seconds. Next, 20 µg of lysate for each sample were loaded onto a 10% BioRad mini-protean TGX gel, along with a Chameleon Duo pre-stained ladder (Licor), and gels were run for 40 minutes at 200 V. Proteins on the gel were transferred to methanol-activated 0.45 µm PVDF membranes in ice-cold transfer buffer (25 mM Tris-HCl at pH 7.6, 192 mM glycine, 20% methanol) at a constant current of 0.35 A for 60 minutes. Membranes were then blocked in a 4% milk solution for 2 hours, incubated in primary antibody solution (monoclonal anti-FLAG M2 from mouse (Sigma) diluted 1:5,000 in 4% milk) for 1 hour at room temperature, and then washed 3 times in TBST buffer for 5 minutes each. Following this wash step, membranes were incubated for 1 hour in secondary antibody solution (goat anti-mouse (Thermo Scientific) diluted 1:5,000 in 4% milk) for another hour at room temperature, then washed 3 more times in TBST for 5 minutes each, followed by 1 wash in TBS for 10 minutes. Finally, 1 mL of working solution from the SuperSignal West Femto Maximum Sensitivity Substrate kit (Thermo Scientific) was added to the membrane. Western blots were imaged using the ‘chemiluminescent+markers’ setting on a Protein Simple FluorChemR imager to visualize bands. Band intensity was quantified by ImageJ to determine relative ratios of proteins.

**β-galactosidase assays**

To measure *rsmY-lacZ* expression, cells (typically ~25 µL) were added to 1 mL of Z buffer (0.06 M $Na_2HPO_4$, 0.04 M $NaH_2PO_4.H_2O$, 0.01 M KCl, 0.001 M $MgSO_4$, adjusted to pH 7, and 135 µL of β-mercaptoethanol added fresh to 50 mL of Z buffer). Next, 50 uL of a 0.1% SDS solution and 100 µL of chloroform were added to the mixture. Each sample was then vortexed for 8 seconds, followed by the addition of 200 µL of a 4 mg/mL ONPG solution. Samples were then vortexed for another 8 seconds and then incubated at 30 $^oC$ for 30 minutes. Following incubation, samples were spun at 21,000 *g* for 2 minutes in a tabletop centrifuge to pellet the chloroform. 250 µL were then withdrawn from the top of each tube and pipetted into a clear bottom 96 well plate to measure both the $OD_{420}$ and $OD_{550}$ of each reaction using a BioTek Synergy H1 plate reader. Miller units were calculated as previously described.

**Protein purification**

To purify proteins for *in vitro* phosphorylation assays, BL21 cells were transformed with the appropriate ML333 or ML310 vectors, in which the expression of MBP-$GacS_c$ or Trx-$RetS_c$ variants are driven by IPTG. Next, 500 mL of each strain was grown up to $OD_{600}$ ~ 0.6 at 37 $^oC$ and induced with 1 mM IPTG for 4 hours at 30 $^oC$. Cells were then spun down for 15 minutes at 5000 *g* and washed twice with lysis buffer (20 mM Tris-HCl, 20 mM imidazole, 0.5 M NaCl, 10% glycerol, 0.1% Triton X-100, adjusted to pH 8). Pellets were resuspended in 10 mL of lysis buffer, with the addition of 1 mg/mL lysozyme, 1 mM PMSF, and 5 µL of benzonase, and incubated for 30 minutes at room temperature. Each sample was lysed by sonication (2 minutes total sonication time, 30 seconds on, 30 seconds off at 30% power). Lysed samples were spun down at 25,000 *g* for 1 hour at 4 degrees to separate the insoluble and soluble fractions. At the same time, Ni-NTA columns (Qiagen) were equilibrated with 10 mL of lysis buffer. Next, the soluble fractions were retained and run through the Ni-NTA columns. Columns were washed with 20 mL of wash buffer (20 mM HEPES, 20 mM imidazole, 0.5 M NaCl, 10% glycerol, 0.1% Triton X-100, adjusted to pH 8). Proteins were then eluted from the column with 2.5 mL of elution buffer (20 mM HEPES, 0.5 M NaCl, 10% glycerol, and 250 mM imidazole, adjusted to pH 8). Eluted proteins were immediately buffer exchanged through PD-10 columns (G.E.), which had been pre-equilibrated with 25 mL of HKEDG storage buffer (10 mM HEPES, 50 mM KCl, 10% glycerol, 0.1 mM EDTA, and 1 mM fresh DTT). Proteins were eluted in 3.5 mL of HKEDG storage buffer from the PD-10 columns and then concentrated using either 10K or 30K MWCO Amicon ultracel centrifugal filters for 30-60 minutes. Protein concentrations were quantified using a Nanodrop spectrophotometer.

***In vitro* phosphorylation assays**

An ATP regeneration system as used for phosphotransfer experiments. GacSΔRecΔHpt was used at a final concentration of 2.5 μM in HKEDG storage buffer. Final concentrations of 0.5 mM ATP, 2.5 μCi of [$\gamma^{32}$P]-ATP, 0.5 mM phosphoenolpyruvate (PEP), and 10 U/mL pyruvate kinase (PK) were first incubated together for 30 minutes at 30 $^{o}$C to allow the $\gamma$-$^{32}$P to equilibrate between ATP and PEP. GacSΔRecΔHpt and $MgCl_2$ were diluted in HKEDG buffer and then added and used at final concentrations of 2.5 μM and 5 mM, respectively. Autophosphorylation reactions were performed at 30 $^{o}$C.

For phosphotransfer assays, GacSΔRecΔHpt was first autophosphorylated using the above parameters at 30 $^{o}$C for 60 minutes, and then spun through a Micro Bio-spin 6 column (Bio-rad) equilibrated with HKEDG buffer to remove unused nucleotide. The cleaned up GacS samples were then incubated with reactions containing RetS and $MgCl_2$ at final concentrations of 5 μM and 5 mM, respectively. Phosphotransfer reactions were run at 30 $^{o}$C. Reactions were stopped at the indicated time points with the addition of 4x SB (200 mM Tris-Cl at pH 6.8, 400 mM DTT, 8% SDS, 0.4% bromophenol blue, 40% glycerol).

Each sample was run on a 10% BioRad mini-protean TGX gel for 50 minutes at 150 V. Gels were then put into zipbloc bags and exposed to phosphor-screens for 4-5 hours so that phosphorylated protein bands could be observed. Screens were imaged using the Typhoon-FLA9500 imager with a “phosphor” setting and a resolution of 50 μm.

**Type VI killing assays**

For killing assays, overnight cultures of PA14 were diluted 1:100 into ABTGC (either with no glycans or 0.1% glycans) and grown for 5 hours at 37 $^{o}$C until reaching an OD of ~0.5.

At the same time, overnight cultures of *E. coli* S17 were diluted 1:500 into ABTGC and grown for 5 hours at 37 $^{o}$C until reaching an OD of ~0.3. 100 µL PA14 and S17 were then spun down at max speed for 2 minutes, and re-suspended in a total volume of 5 µL to concentrate the cells. This suspension was then plated on ABTGC plates (3% agar), and incubated at 37 $^{o}$C for 1.5 hours. The mixture was then scraped off the plates and re-suspended in 100 µL of ABTGC, and then serially diluted by 10x to count cfus. These serially diluted samples were plated on either LB-irgasan plates (to select for PA14) or LB-cefsulodin plates (to select for S17) overnight at 37 $^{o}$C. Irgasan and cefsulodin were used at a concentration of 25 µg/mL. Competitive index was calculated as: CI = $E_o$ / $E_t$ / $P_t$, where E is the number of *E. coli* cells competed with or without *P. aeruginosa* ($E_t$ and $E_0$, respectively) and $P_t$ is the number of competing *P. aeruginosa* cells.

### Supplementary Figures

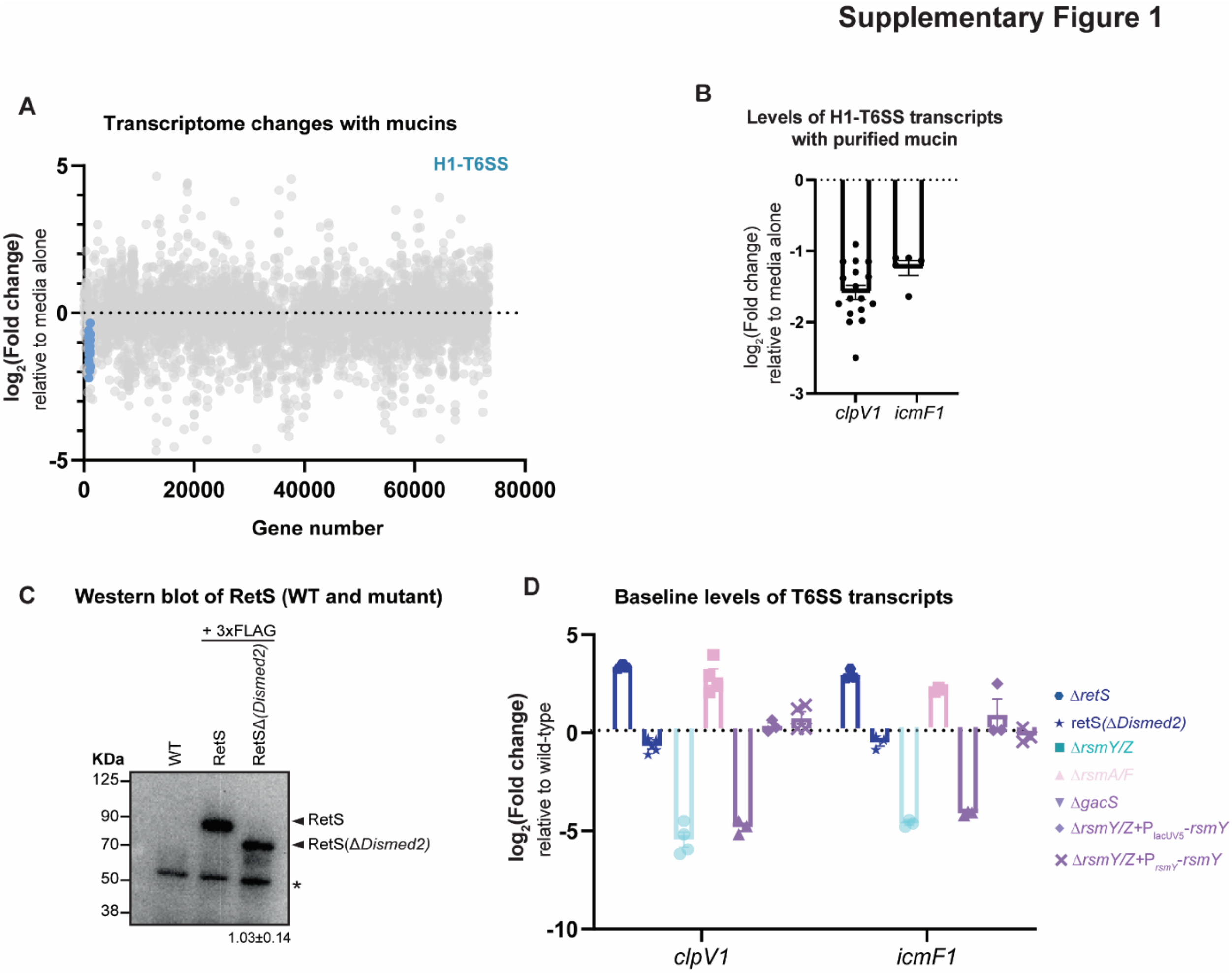


**Figure S1: Gene expression analysis of differentially regulated genes in response to mucin in the wild type, and baseline levels of H1-T6SS transcripts in different mutant backgrounds.**

A) RNA-seq analysis of gene expression changes in response to 0.5% mucin, relative to a media control. RNA-seq data are plotted as a function of genomic position (denoted by gene number), with each circle representing an individual gene. The H1-T6SS operon is shown with blue circles.

B) Levels of T6SS transcripts in following exposure to isolated mucin glycoproteins relative to medium alone. Gene expression measured by qRT-PCR and normalized to a control gene (*rpoD*). Bars indicate the mean ± SEM, with individual measurements shown.

C) Representative Western blot of WT RetS-3xFLAG and RetS(Δ*Dismed2*)-3xFLAG. The quantification (mean ± SD, n=4) of RetS(Δ*Dismed2*)-3xFLAG levels relative to WT RetS-3xFLAG levels is indicated below. * indicates a non-specific band also present in the WT strain.

D) Baseline levels of *clpV1* and *icmF1* in each GacS/Rsm/RetS mutant relative to the WT. Gene expression measured by qRT-PCR and normalized to a control gene (*rpoD*). Bars indicate the mean ± SEM, with individual measurements shown (dots).

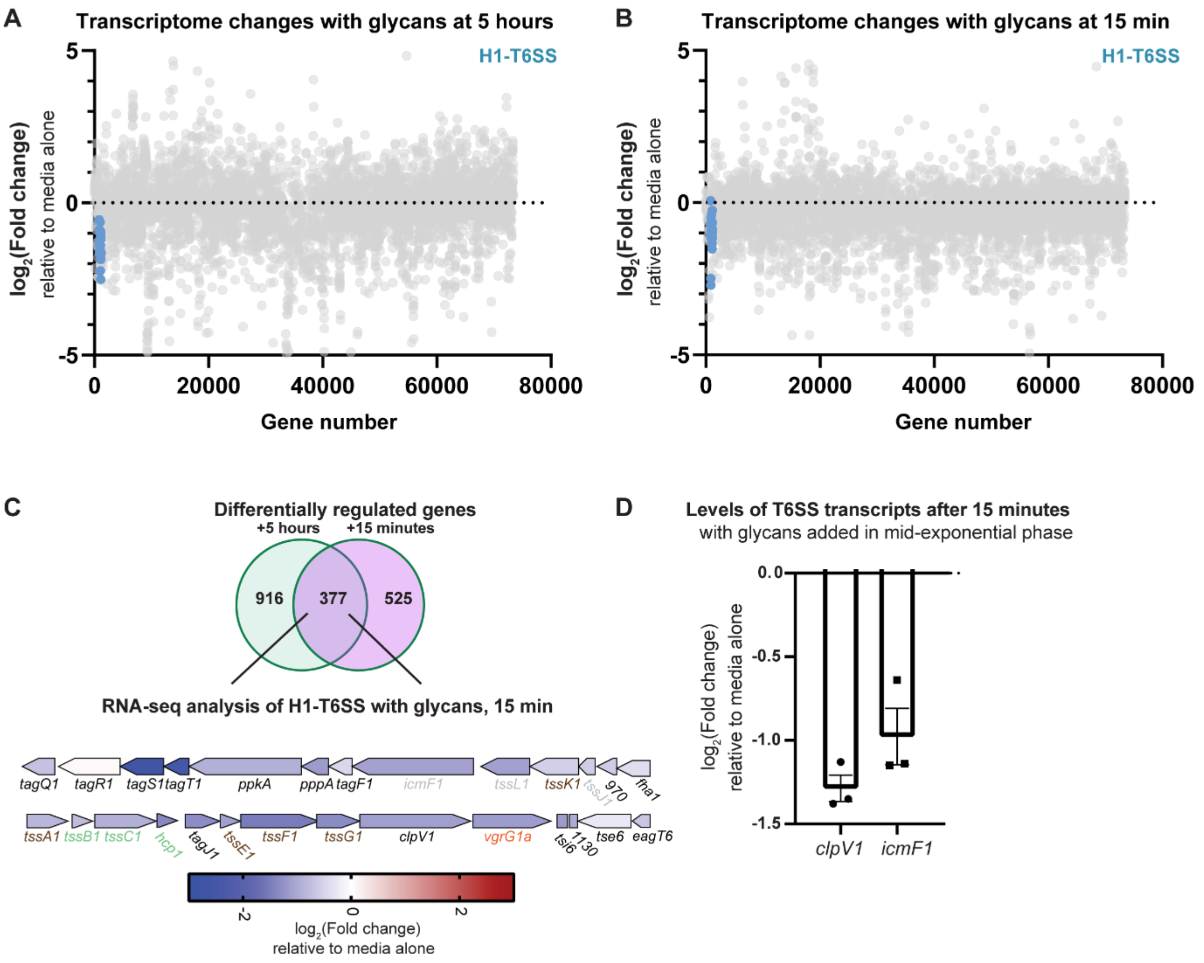


**Figure S2: Mucin glycans downregulate the H1-T6SS at both 5 hours and 15 minutes after exposure.**

A) RNA-seq analysis of gene expression changes in response to 0.1% glycans after 5 hours of exposure, relative to a media control. RNA-seq data are plotted as a function of genomic position (denoted by gene number), with each circle representing an individual gene. The H1-T6SS operon is shown with blue circles.

B) RNA-seq analysis of gene expression changes in response to 0.1% glycans after 15 minutes of exposure, relative to a media control. RNA-seq data are plotted as a function of genomic position (denoted by gene number), with each circle representing an individual gene. The H1-T6SS operon is shown with blue circles.

C) Overlap of differentially expressed genes following glycan exposure for 5 hours or 15 minutes relative to medium alone, measured by RNA-sequencing. Diagram of the H1-T6SS operon representing the fold change of each gene following exposure to glycans for 15 minutes relative to medium alone, measured by RNA-sequencing. Arrows indicate the orientation and relative size of each gene. The names of genes that encode key components of the T6SS apparatus are color coded according to Fig. 1B (baseplate in brown, membrane complex in gray, needle/sheath in green, tip in orange).

D) Levels of T6SS transcripts in PA14 following exposure to the pool of mucin glycans for 15 minutes relative to medium alone. Gene expression measured by qRT-PCR and normalized to a control gene (*rpoD*). Bars indicate the mean ± SEM, with individual measurements shown (black dots).

### Supplementary Figure 3

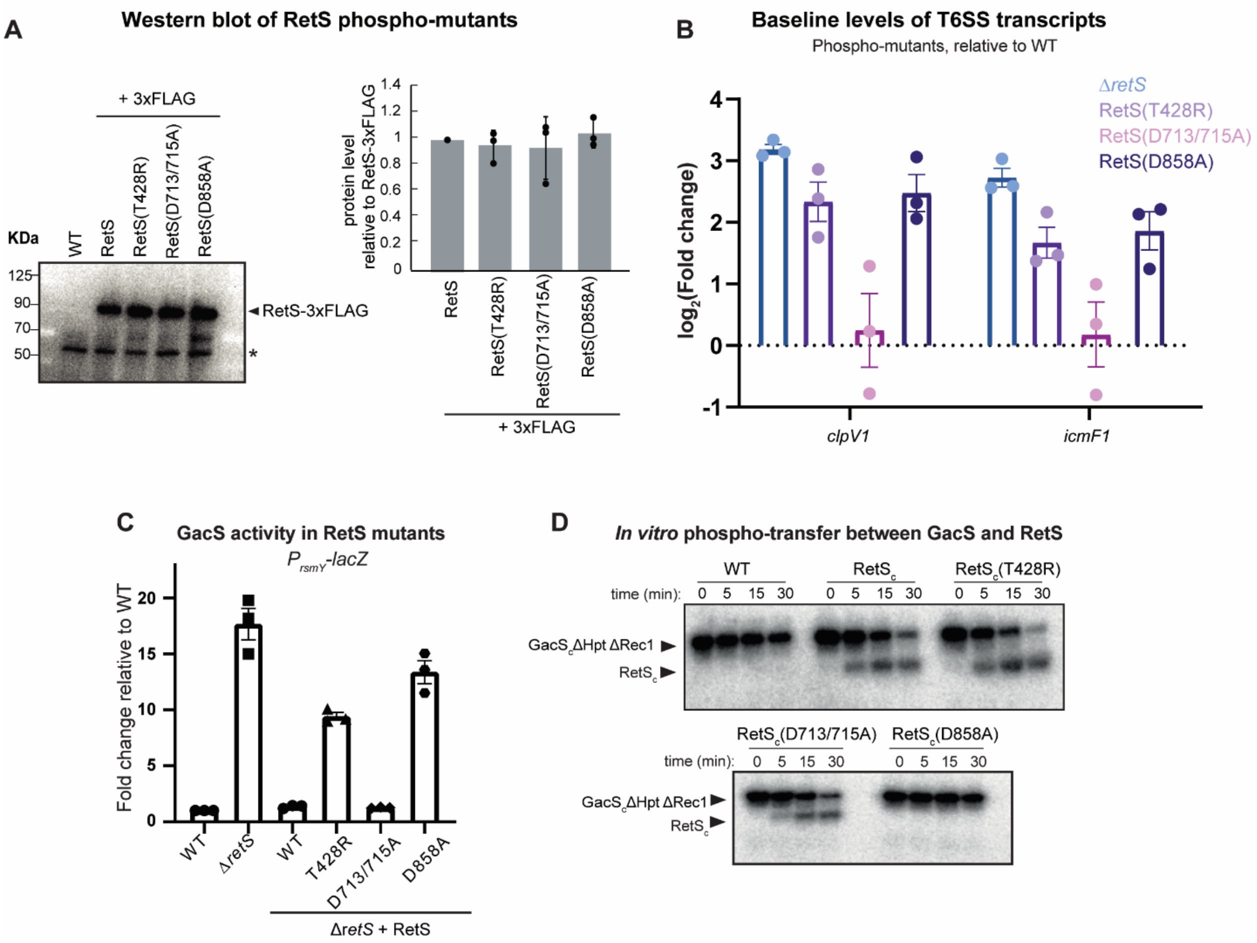


**Figure 3: Measurement of GacS activity and RetS stability in RetS phosphor-mutants.**

A) Western blot of 3x-FLAG tagged versions of chromosomal RetS point mutants. A quantification from three independent blots is shown on the right, indicating mean ± SD, with individual values indicated (black dots). * indicates a non-specific band also present in the WT strain.

B) Baseline levels of *clpV1* and *icmF1* in each RetS mutant relative to the WT. Gene expression measured by qRT-PCR and normalized to a control gene (*rpoD*). Bars indicate the mean ±SEM, with individual measurements shown (dots).

C) *rsmY-lacZ* expression, quantified by β-galactosidase assays, in WT and Δ*retS* strains and a series of strains in which *retS* was deleted and then replaced, at the native locus, with the mutant variant of *retS* indicated (or a wild-type control). Data shown are mean ± SEM, with individual values indicated.

D) Phosphotransfer from $GacS_c$ΔRec1ΔHpt (the GacS kinase core) to $RetS_c$. $GacS_c$ΔRec1ΔHpt was autophosphorylated and then examined for phosphotransfer to wild-type $RetS_c$, the $RetS_c$ point mutants indicated, or a buffer control. Reaction times (in minutes) are indicated above each gel. Bands corresponding to various phosphorylated proteins are labeled on the left of each gel. GacS is able to phosphorylate RetS, except in the RetS(D858A) mutant, indicating that the second receiver domain of RetS is responsible for siphoning phosphate from GacS.

### Supplementary Figure 4

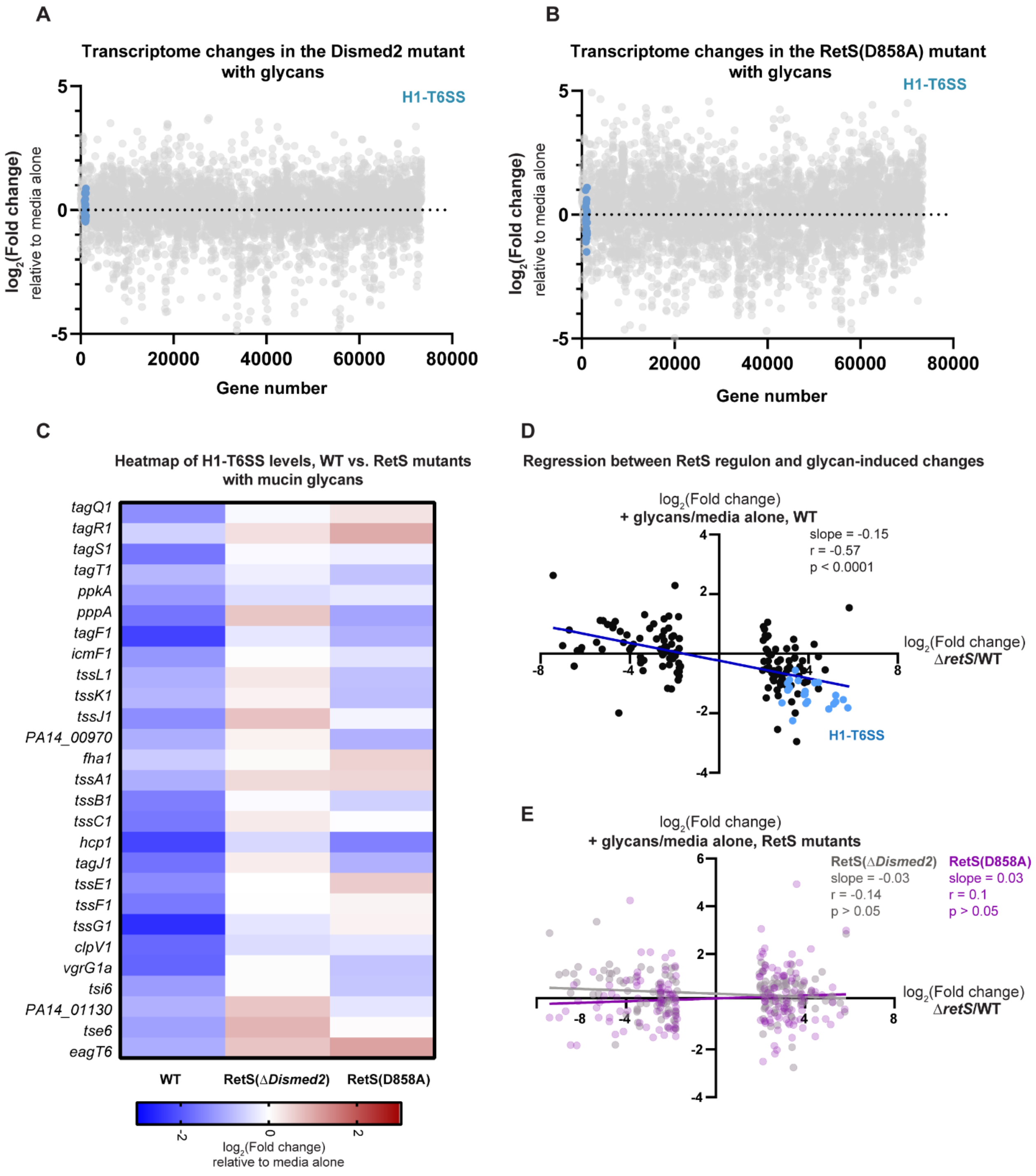


**Figure S4: Mucin glycans differentially regulate the RetS regulon in a RetS-dependent manner.**

A) RNA-seq analysis of gene expression changes in response to 0.1% glycans in the RetS(Δ*Dismed2*) strain 5 hours after exposure, relative to a media control. RNA-seq data are plotted as a function of genomic position (denoted by gene number), with each circle representing an individual gene. The H1-T6SS operon is shown with blue circles.

B) RNA-seq analysis of gene expression changes in response to 0.1% glycans in the RetS(D858A) strain 5 hours after exposure, relative to a media control. RNA-seq data are plotted as a function of genomic position (denoted by gene number), with each circle representing an individual gene. The H1-T6SS operon is shown with blue circles.

C) Heatmap of all H1-T6SS transcript levels following exposure to mucin glycans for 5 hours relative to a medium control in the wild type, RetS(Δ*Dismed2*), and RetS(D858A) strains, as measured by RNA-sequencing.

D,E) Regression of glycan-induced transcript-level changes in the wild-type (D) or RetS mutants (E) against the top 50% of differentially expressed genes in the RetS regulon (Δ*retS/*WT) (*12*), that were also present and expressed in the PA14 RNA-seq datasets. Genome-wide transcriptional changes were measured by RNA-sequencing. H1-T6SS genes are highlighted in blue in the top panel. In the bottom panel, RetS(Δ*Dismed2*) is highlighted in gray, and RetS(D858A) in purple.

### Supplementary Figure 5

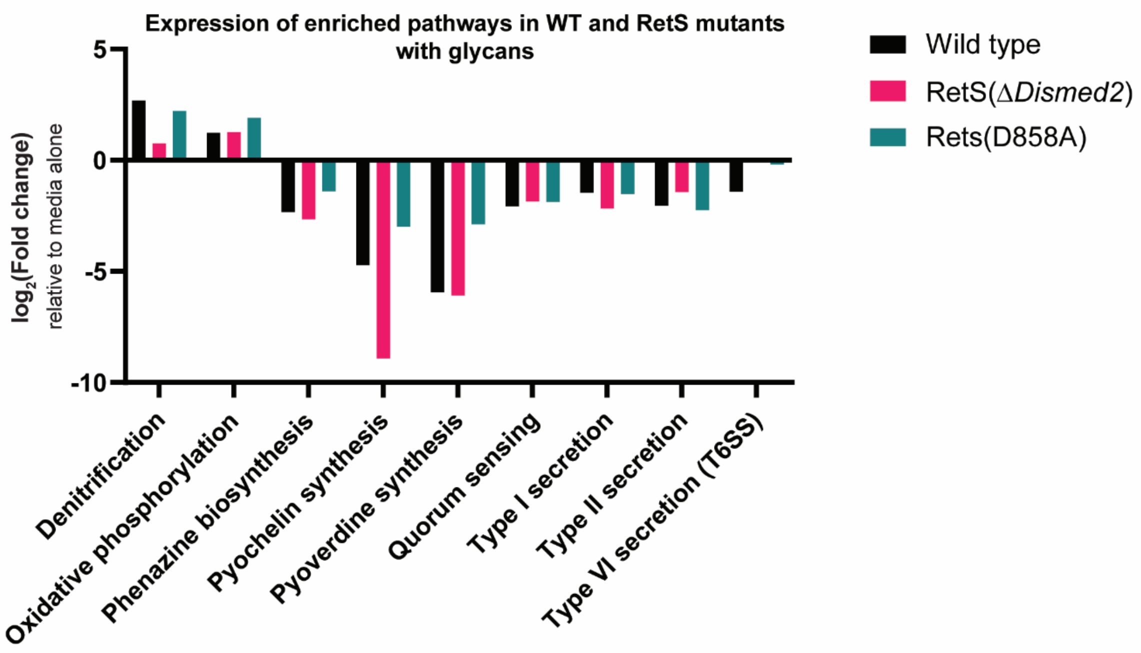


**Figure S5: Mucin glycans regulate other pathways independently of RetS.**

Average expression of each enriched pathway upon exposure to 0.1% glycans in the WT (black), RetS(Δ*Dismed2*) (red), and RetS(D858A) (green) strains. Fold changes are derived from each respective RNA-seq dataset.

**Supplementary Figure 6**

A

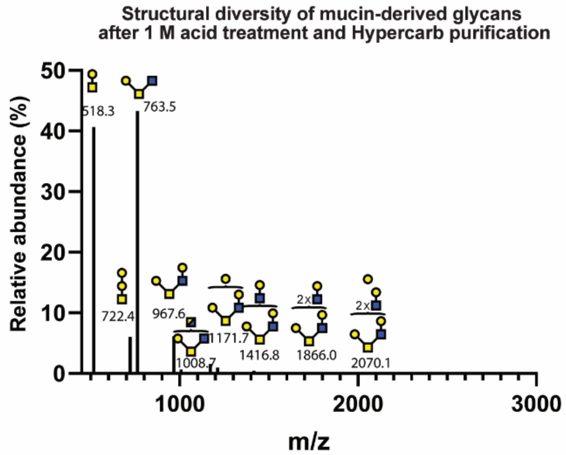


B

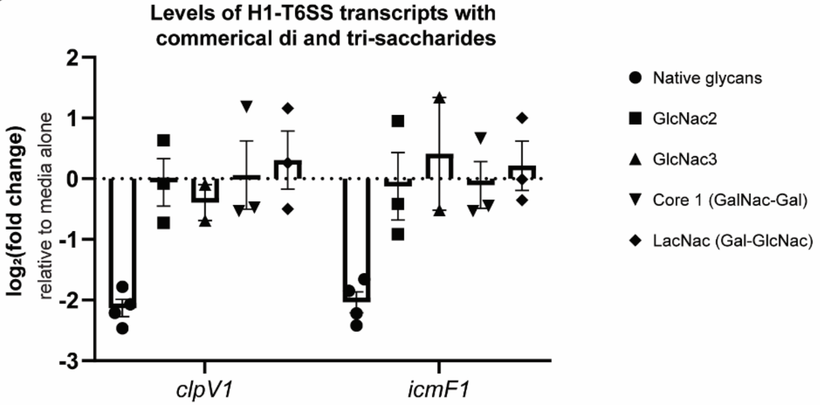


**Figure S6: Acid-treated mucin glycans are retained on Hypercarb columns, and commercially available synthetic moieties present on mucin glycans do not downregulate H1-T6SS transcripts.**

A) MALDI-TOF-MS spectrum of MUC5AC glycans following 1 M TFA treatment and Hypercarb purification. Selected peaks labeled with monoisotopic masses and predicted structures.

B) Levels of T6SS transcripts in PA14 following exposure to commercially available di and tri-saccharides. Gene expression measured by qRT-PCR and normalized to a control gene (*rpoD*). Bars indicate the mean ± SEM, with individual measurements shown (black dots).

### Supplemental tables 1-3 are excel spreadsheets (due to size)

### Table S4: *Pseudomonas aeruginosa* strains

| Strain | Description | Reference |
|---|---|---|
| **ML3006** | WT PA14 | lab collection |
| **ML3007** | Δ*retS* | This study |
| **ML3008** | Δ*gacS* | This study |
| **ML3009** | WT PA14 + *rsmY-lacZ* | This study |
| **ML3012** | RetS(Δ*Dismed2)* + *rsmY-lacZ* | This study |
| **ML3013** | RetS(Δ*Dismed2)* + -3x-FLAG + *rsmY-lacZ* | This study |
| **ML3015** | RetS-3x-FLAG + *rsmY-lacZ* | This study |
| **ML3016** | Δ*retS* + RetS | This study |
| **ML3017** | Δ*retS* + RetS(T428R) | This study |
| **ML3018** | Δ*retS* + RetS(D713/715A) | This study |
| **ML3019** | Δ*retS* + RetS(D858A) | This study |
| **ML3020** | Δ*retS* + RetS + *rsmY-lacZ* | This study |
| **ML3021** | Δ*retS* + RetS(T428R) + *rsmY-lacZ* | This study |
| **ML3022** | Δ*retS* + RetS(D713/715A) + *rsmY-lacZ* | This study |
| **ML3023** | Δ*retS* + RetS(D858A) + *rsmY-lacZ* | This study |
| **ML3024** | Δ*retS* + RetS(T428R)-3xFLAG + *rsmY-lacZ* | This study |
| **ML3025** | Δ*retS* + RetS(D713/715A)-3xFLAG + *rsmY-lacZ* | This study |
| **ML3026** | Δ*retS* + RetS(D858A)-3xFLAG + *rsmY-lacZ* | This study |
| **ML3249** | Δ*rsmY/Z* | This study |
| **ML3250** | Δ*rsmA/F* | This study |
| **ML3251** | Δ*rsmY/Z*+$P_{lacUV5}$-*rsmY* | This study |
| **ML3252** | Δ*rsmY/Z*+$P_{rsmY}$-*rsmY* | This study |
| **ML3253** | Δ*icmF1* | This study |

### Table S5: *Pseudomonas aeruginosa* plasmids

| Strain | Plasmid | Description | Reference |
|---|---|---|---|
| **ML2827** | **Mini-CTX-lacZ** | For chromosomal integration of genes in *P. aeruginosa* | (*31*) |
| **ML2828** | **pCTX-*rsmY-lacZ*** | Chromosomal integration of *rsmY-lacZ* | (*32*) |
| **ML2829** | **pexG2** | Suicide vector for allelic exchange | (*33*) |
| **ML2830** | **pMQ30** | Suicide vector for allelic exchange | (*34*) |
| **ML3263** | **pPSV38** | Plasmid-borne expression of genes | (*35*) |
| **ML2831** | **pEXG2-RetS(Δ*Dismed2)*** | Chromosomal deletion of Dismed2 domain of RetS | This study |
| **ML2832** | **pEXG2-RetS-3xFLAG** | Chromosomal integration of 3x-FLAG tag onto the C terminus of RetS | This study |
| **ML2833** | **pEXG2-RetS(D858A)-3xFLAG** | Chromosomal integration of 3x-FLAG onto the C terminus of RetS(D858A) mutant | This study |
| **ML2835** | **pMQ30-RetS(T428R)** | Integrate mutant RetS at native RetS locus on chromosome | This study |
| **ML2836** | **pMQ30-RetS(D713/715A)** | Integrate mutant RetS at native RetS locus on chromosome | This study |
| **ML2837** | **pMQ30-RetS(D858A)** | Integrate mutant RetS at native RetS locus on chromosome | This study |
| **ML3254** | **pEXG2-*rsmY*** | Chromosomal deletion of *rsmY* | This study |
| **ML3255** | **pEXG2-*rsmZ*** | Chromosomal deletion of *rsmZ* | This study |
| **ML3256** | **pEXG2-*rsmA*** | Chromosomal deletion of *rsmA* | This study |
| **ML3257** | **pEXG2-*rsmF*** | Chromosomal deletion of *rsmF* | This study |

| | | | |
|---|---|---|---|
| **ML3258** | **pEXG2-*icmF1*** | Chromosomal deletion of *icmF1* | This study |
| **ML3259** | **pPSV38-** $_{lacUV5}$**-*rsmY*** | Express *rsmY* from constitutive lacUV5 promoter | This study |
| **ML3260** | **pPSV38-** $_{rsmY}$**-*rsmY*** | Express *rsmY* from native *rsmY* promoter | This study |
| **ML3261** | **pMQ30-GacS** | Chromosomal deletion of GacS | This study |
| **ML3262** | **pMQ30-RetS** | Chromosomal deletion of RetS | This study |

### Table S6: *E. coli* expression plasmids

| Strain | Plasmid | Description | Reference |
|---|---|---|---|
| **ML310** | ML310 | IPTG-driven expression of Trx-tagged proteins | Lab collection |
| **ML333** | ML333 | IPTG-driven expression of MBP-tagged proteins | Lab collection |
| **ML2840** | ML333-GacSΔHpt ΔRec1 | Expression of $GacS_c$ DHp/CA only | This study |
| **ML2842** | ML310-RetS | Expression of $RetS_c$ | This study |
| **ML2843** | ML310-RetS(T428R) | Expression of putatively phosphatase dead RetS | This study |
| **ML2844** | ML310-RetS(D713/715A) | Expression of receiver one dead RetS | This study |
| **ML2845** | ML310-RetS(D858A) | Expression of receiver two dead RetS | This study |

### Table S7: Primers

| # | Description | Sequence |
|---|---|---|
| **1** | pEXG2_vector_ΔDISMED2_F | GGCTGATCCAGCAGCTCAACCTGCAACAGCTGCTTTACATTTATG CTTCCGGCTCGTA |
| **2** | pEXG2_vector_ΔDISMED2_R | AGGACCAGGGAGGACTCCAGGCGGACCATGCTTAATTAATTTCC ACGGGTGCGCATG |
| **3** | pEXG2_ DISMED2_frag1_F | GATCATGCGCACCCGTGGAAATTAATTAAGCATGGTCCGCCTGG AGTCCTCCCTGGTCCT |
| **4** | pEXG2_ DISMED2_frag1_R | GAGCAGCATGCCGAAGGCGTAGGCGGGCTTGGCGCTGGGAGTA GTGGCGGTGGTTTGCAG |
| **5** | pEXG2_ DISMED2_frag2_F | CTGCAAACCACCGCCACTACTCCCAGCGCCAAGCCCGCCTACGCC TTCGGCATGCTGCTC |
| **6** | pEXG2_ DISMED2_frag2_R | TATACGAGCCGGAAGCATAAATGTAAAGCAGCTGTTGCAGGTTG AGCTGCTGGATCAGCC |
| **7** | pEXG2-RetS-3xFLAG_vec_F | CGGCCGCTACATCTTCATCCTCTTCGGTATTGCTTTACATTTATGC TTCCGGCTCGTATA |
| **8** | pEXG2-RetS-3xFLAG_vec_R | TCGCCGCCAGTTGCATGCCGGTCATGCCGCTTAATTAATTTCCAC GGGTGCGCATGATC |
| **9** | pEXG2-RetS-3xFLAG_insert_F | GATCATGCGCACCCGTGGAAATTAATTAAGCGGCATGACCGGCA TGCAACTGGCGGCGA |
| **10** | pEXG2-RetS-3xFLAG_insert_R | TATACGAGCCGGAAGCATAAATGTAAAGCAATACCGAAGAGGA TGAAGATGTAGCGGCCG |
| **11** | pEXG2-RetS-3xFLAG_RTH_F | AGATCATGACATCGACTACAAGGATGACGATGACAAGTGAGGG CAGCGACGTGCTCCGGC |
| **12** | pEXG2-RetS-3xFLAG_RTH_R | TTATAATCACCGTCATGGTCTTTGTAGTCGCCGCTACCGCCGGAG GGCAGGGCGTCGCCC |
| **13** | pEXG2-GacS-3xFLAG_vec_F | GAAACCCGGCGCGATGCTGATCAATACCGGTGCTTTACATTTATG CTTCCGGCTCGTATA |
| **14** | pEXG2-GacS-3xFLAG_vec_R | CGACGCAGAGCAGCCGTGGCGGCCGTCCGGCTTAATTAATTTCC ACGGGTGCGCATGATC |
| **15** | pEXG2-GacS-3xFLAG_insert_F | GATCATGCGCACCCGTGGAAATTAATTAAGCCGGACGGCCGCCA CGGCTGCTCTGCGTCG |
| **16** | pEXG2-GacS-3xFLAG_insert_R | TATACGAGCCGGAAGCATAAATGTAAAGCACCGGTATTGATCAG CATCGCGCCGGGTTTC |

| | | |
|---|---|---|
| **17** | pEXG2-GacS-3xFLAG_RTH_F | AGATCATGACATCGACTACAAGGATGACGATGACAAGTGACCATGCGCATCCTGTTCTTC |
| **18** | pEXG2-GacS-3xFLAG_RTH_R | TTATAATCACCGTCATGGTCTTTGTAGTCGCCGCTACCGCCGAGTTCGCTGGAGTCGAGG |
| **19** | pEXG2-RetS(D858A)-3xFLAG-RTH_F | GCCTGCGAGATGCCGGTGCTCGACGGC |
| **20** | pEXG2-RetS(D858A)-3xFLAG-RTH_R | CATCAGCACCAGGTCGTACTGGGTC |
| **21** | RetS(T428R)_F | ACGAGATCCGCAGGCCCATGAACGGCG |
| **22** | RetS(T428R)_R | CGCCGTTCATGGGCCTGCGGATCTCGT |
| **23** | RetS(D713/715A)_F | CCTGCTCGCCCAGGCCATGCCCGG |
| **24** | RetS(D713/715A)_R | CCGGGCATGGCCTGGGCGAGCAGG |
| **25** | RetS(D858A)_F | GGTGCTGATGGCCTGCGAGATGCCGGT |
| **26** | RetS(D858A)_R | ACCGGCATCTCGCAGGCCATCAGCACC |
| **27** | pENTR-RetS_F | caccATCCAGCAGCTCAACCTGCAACAGCGCAC |
| **28** | pENTR-RetS_R | TCAGGAGGGCAGGGCGTCGCCCTGG |
| **29** | pENTR-GacS_F | caccGGCAGCAACGAGCTGGACGAACTGGCCTCC |
| **30** | pENTR-GacS_R | TCAGAGTTCGCTGGAGTCGAGGCTGGTGGAA |
| **31** | pENTR-GacS ΔRec1 ΔHpt_R | TCATGGCGGCCGTCCGGAAACCATGGCGTG |
| **32** | Δ*retS*_primer1 | caggctgaaaatcttctctcatccgccaaaGCCTACCTGCGCGAGCAGGGCGC |
| **33** | Δ*retS*_primer2 | TCGCCCTGGCGGCGGCGATCGATCCGAAGCCGTACCACGGCGAAGTCCCTTC |

| | | |
|---|---|---|
| **34** | Δ*retS*_primer3 | CCGTGGTACGGCTTCGGATCGATCGCCGCCGCCAGGGCGACG |
| **35** | Δ*retS*_primer4 | GCGGATAACAATTTCACACAGGAAACAGCTCGCCAGCGCGCAGACGAACAGACCC |
| **35** | Δ*gacS*_primer1 | caggctgaaaatcttctctcatccgccaaaACTACGTCCCCAGGTCCAGGACAGCCATACCGATAG |
| **36** | Δ*gacS*_primer2 | TCG AGG CTG GTG GAA GAC AGG CCG AGA TCC TTG AAC ACA CGT CTC TCC GTC G |
| **38** | Δ*gacS*_primer3 | GTGTGTTCAAGGATCTCGGCCTGTCTTCCACCAGCCTCGACTCCAGCGAAC |
| **39** | Δ*gacS*_primer4 | GCG GAT AAC AAT TTC ACA CAG GAA ACA GCT TGC TAC CAC GCC ATC AGG CCC GGA CCC G |
| **40** | Δ*rsmY*_primer1 | GATCATGCGCACCCGTGGAAATTAATTAAGCGGTGGCCACGTAGTTCGGGG |
| **42** | Δ*rsmY*_primer2 | CGACGCGGTTTTCCTCGGGCAATAAGGTTTGAAGATTACGCATCTCTGCGAGGG |
| **43** | Δ*rsmY*_primer3 | CCCTCGCAGAGATGCGTAATCTTCAAACCTTATTGCCCGAGGAAAACCGCGTCG |
| **44** | Δ*rsmY*_primer4 | TATACGAGCCGGAAGCATAAATGTAAAGCACTGCTCACCGGCAACCTGGATATCG |
| **45** | Δ*rsmZ*_primer1 | GATCATGCGCACCCGTGGAAATTAATTAAGCATGCTCGGCCTGAACGAACGGG |
| **46** | Δ*rsmZ*_primer2 | GGCGACGAGTAAAACGGCAGGCAAACGCAGGAGTGATATTAGCGATTCCCTG |
| **47** | Δ*rsmZ*_primer3 | CAGGGAATCGCTAATATCACTCCTGCGTTTGCCTGCCGTTTTACTCGTCGCC |
| **48** | Δ*rsmZ*_primer4 | TATACGAGCCGGAAGCATAAATGTAAAGCAGCACGAGATGCCGAGCCAGCAG |
| **49** | Δ*rsmA*_primer1 | GATCATGCGCACCCGTGGAAATTAATTAAGCCTTCAAGATCCTCGGGCCGATC |
| **50** | Δ*rsmA*_primer2 | TCGGCGCGTTGACGCCGAAATCAGCATTCCTTTCTCCTCACGCGAATATTTC |
| **51** | Δ*rsmA*_primer3 | GAAATATTCGCGTGAGGAGAAAGGAATGCTGATTTCGGCGTCAACGCGCCGA |

| | | |
|---|---|---|
| **52** | Δ*rsmA*_primer4 | TATACGAGCCGGAAGCATAAATGTAAAGCAGCAACTGTCGATCCTTCGTCCGTC |
| **53** | Δ*rsmF*_primer1 | GATCATGCGCACCCGTGGAAATTAATTAAGGCTCCAGGTTGAGCTGATTGAGGC |
| **54** | Δ*rsmF*_primer2 | CTTTCGGTGCCGTCTTCAACTCGTCGAAACCCATGTTCCGCGTCCTTGC |
| **55** | Δ*rsmF*_primer3 | GCAAGGACGCGGAACATGGGTTTCGACGAGTTGAAGACGGCACCGAAAG |
| **56** | Δ*rsmF*_primer4 | TATACGAGCCGGAAGCATAAATGTAAAGCATAATCGCGTTCGGCCTGCTGG |
| **57** | Δ*icmF1*_primer1 | GATCATGCGCACCCGTGGAAATTAATTAAGATCGCCGACGCCCTGCGCAAGGTCAAGG |
| **58** | Δ*icmF1*_primer2 | GGCCAGCTTGCCGTAGAAACCGACGCTGTGACCTTCGCCGCGTTGCGCCGGGCCT |
| **59** | Δ*icmF1*_primer3 | AGGCCCGGCGCAACGCGGCGAAGGTCACAGCGTCGGTTTCTACGGCAAGCTGGCC |
| **60** | Δ*icmF1*_primer4 | TATACGAGCCGGAAGCATAAATGTAAAGCAGCGCTGCGCCGGGGTCACCGCTTC |
| **61** | $P_{lacUV5}$-*rsmY* primer 1 | GCTTCCGGCTCGTATAATGTGTGGGTCAGGACATTGCGCAGGAAG |
| **62** | $P_{lacUV5}$-*rsmY* primer 2 | GTTTAGAGGCCCCAAGGGGTTATGCTAAAAGGCGTGGTCTGAGCGAC |
| **63** | $P_{rsmY}$-*rsmY* primer 1 | CGGTACCCGGGGATCCTCTAGAGCGAGCGGAACTATTACAGCGTGT |
| **64** | $P_{rsmY}$-*rsmY* primer 2 | GTTTAGAGGCCCCAAGGGGTTATGCTAAAAGGCGTGGTCTGAGCGACG |
| **65** | *clpV1*_F (qPCR) | CACAAGGTGCCGTTCGAGTT |
| **66** | *clpV1*_R (qPCR) | GTTGTCGCGGCTACTGATAC |
| **67** | *icmF1*_F (qPCR) | CAACCCTACGTCGACACCTC |
| **68** | *icmF1*_R (qPCR) | CATGGTCACCGGCTTGAGTT |

| | | |
|---|---|---|
| **69** | *rpoD*_F (qPCR) | GGGCGAAGAAGGAAATGGTC |
| **70** | *rpoD*_R (qPCR) | CAGGTGGCGTAGGTAGAGAA |

**Tables S1 to S3 titles**

Table S1: RNA-seq data for all conditions (+/- mucin, +/- glycans, and in each strain)

Table S2: All MS data for glycan analysis.

Table S3: Source data for regressions in Fig S4D-E.
